# Extracting large-scale neural activity with Suite2p

**DOI:** 10.64898/2026.02.04.703741

**Authors:** Carsen Stringer, Chris Ki, Nicholas DelGrosso, Paul LaFosse, Qingqing Zhang, Lin Zhong, Marius Pachitariu

## Abstract

Neural recordings using optical methods have been steadily improving. For example, we demonstrate here recordings of over 100,000 neurons from the mouse cortex obtained with a standard commercial microscope. To process such large datasets, we developed Suite2p, a collection of efficient algorithms for motion correction, cell detection, activity extraction and quality control. We also developed new approaches to benchmark performance on these tasks. Our GPU-accelerated non-rigid motion correction outperforms alternative methods, while running over five times faster. For cell detection, Suite2p outperforms the CNMF algorithm in Caiman and Fiola, finding more cells and producing fewer false positives, while running in a fraction of the time. We also introduce quality control steps for users to evaluate performance on their own data, while offering alternative algorithms for specialized types of recordings such as those from one-photon imaging.

## Introduction

Optical recording methods are useful in a wide variety of neuroscience experiments [1, 2]. They allow for easy multiplexing of an activity indicator with other optical markers, for example to determine cell types or projection targets [3]; they give exact tissue position for each recorded neuron, and enable sampling of wide areas to look for changes in tuning within and across brain areas [4, 5]; they allow for easy demixing of activity between nearby neurons simply in virtue of their spatial separation; they enable tracking of the activity of large neural populations across weeks and months [6, 7]; they also allow for scaling up of neuronal yields at the expense of temporal resolution [8, 9]; miniaturized microscopes enable imaging of the same population over many hours [10, 11] etc. However, optical recordings typically produce large amounts of raw data, which need to be analyzed and processed in order to obtain interpretable neural activity timecourses. Suite2p is a collection of algorithms developed specifically for this purpose.

In the analysis of optical recordings, two processing steps have attracted the most research attention, namely the spike detection step and the cell detection step. For spike detection, we do not introduce a new algorithm here and instead rely on a fast implementation of OASIS [12] (see Discussion). Cell detection is the task of creating pixel masks that cover the somas or other biological compartments of a cell. This is typically done using functional data, because pixels in different cells are high at different times, while pixels in the same cell are active simultaneously [13–25]. The task of cell detection is well-matched to matrix decomposition problems, for which a large number of approaches are readily available. The Suite2p detection algorithm can also be interpreted as a matrix decomposition approach, but it relies on strong L0 sparsity constraints on both matrix factors, for which standard solutions do not exist.

Less research attention has been directed towards the registration and quality control steps. While most types of registration rely on frame correlation approaches, the implementation and accuracy of these methods can vary substantially, as we shall see below. Recent methods have claimed improvements in processing speed from running on GPUs, though we have found these claims to only be valid at very small frame sizes (*<*200 pixels). Finally, some type of quality control step would be useful after each data processing step, but there has been limited development in this area. For example, quality control steps can show a user whether the registration was successful, whether Z drift was encountered in the recording, whether cells have been extracted uniformly from across the entire field of view, whether neural activity patterns are dominated by artifacts, etc.

## Results

The development of Suite2p has continued uninterrupted over the past 10 years, starting with the algorithms in the original preprint [27]. Here we describe and evaluate the algorithms in Suite2p as they exist today, serving as a reference at this particular point in time. The current version (v1.0) (Figure 1a) includes: 1) support for multiple input file types; 2) GPU-accelerated motion registration in 2D and across Z-stacks; 3) multiple algorithms for cell segmentation relying on either functional or anatomical information; 4) GPU-accelerated activity extraction with neuropil correction and spike deconvolution; and 5) ROI classification and quality control, integrated in a graphical user interface (GUI) with extensive functionality (Figure 1b, Figure S1). Suite2p completes all tasks in a fraction of the acquisition time, for two-photon imaging, on a standard desktop computer with a consumer-grade GPU. The software package is available at github.com/MouseLand/suite2p.

**Figure 1:**
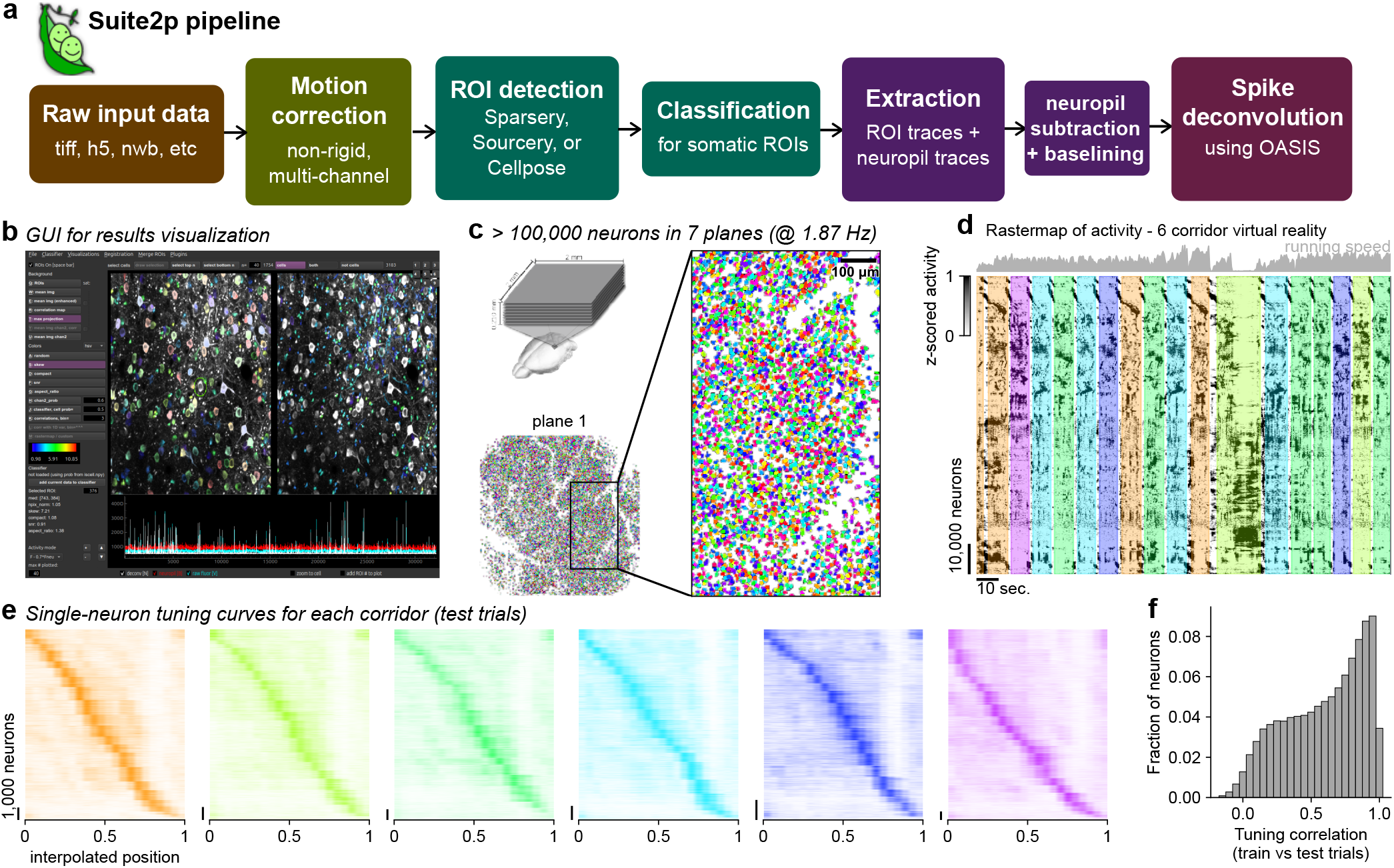
Suite2p pipeline. **a**, Outline of the Suite2p pipeline. **b**, The graphical user interface (GUI) can be used for manual curation and results visualization. **c**, Multi-plane recording of over 100,000 neurons using a standard Thorlabs Bergamo2 scope (see Methods for exact configuration). Somatic ROIs for example plane shown in different colors. **d**, Neural activity raster from recording in **c**, with neurons sorted by Rastermap, colored by the corridor that the mouse is in [26]. **e**, Single-neuron tuning curves where each neuron is plotted at its preferred position in the corridor in which it responded highest. Preferred position and corridor were chosen using held-out (odd) trials, and tuning curves are shown on test (even) trials. **f**, Per-neuron correlation of tuning curves between train and test trials, across all corridors.

We demonstrated in the original Suite2p preprint that large-scale recordings of over 10,000 neurons are possible on commercially-available microscopes such as the Thorlabs Bergamo2, which served as motivation for the development of highly-efficient algorithms that are also highly-accurate so that manual curation was no longer necessary. Here we show that on the same type of microscope, recording size can be pushed above 100,000 neurons simultaneously, further emphasizing the requirements of high efficiency and high accuracy for data analysis (Figure 1c and Figure S2a). Due to the relatively large spacing between planes (30*µ*m) and the precise optical sectioning of the microscope, the ∼100,000 neurons are mostly unique (Figure S2bc), with no relation between the tuning curves of each neuron and those of neurons at the same location in the next plane (Figure S2d). Datasets at this scale allow for large-scale population analyses and visualizations such as Rastermap (Figure 1d), while providing dense sampling of neurons with strong tuning to many sensory (Figure 1ef) or motor features. The increase of recording scale was made possible by technological upgrades to the microscope, a new transgenic mouse line [28], and improvements of the Suite2p detection algorithm for smaller ROIs. We note that Suite2p can also be used for recordings at standard zoom and imaging rates (i.e. Figure 2 and Figure 4).

**Figure 2:**
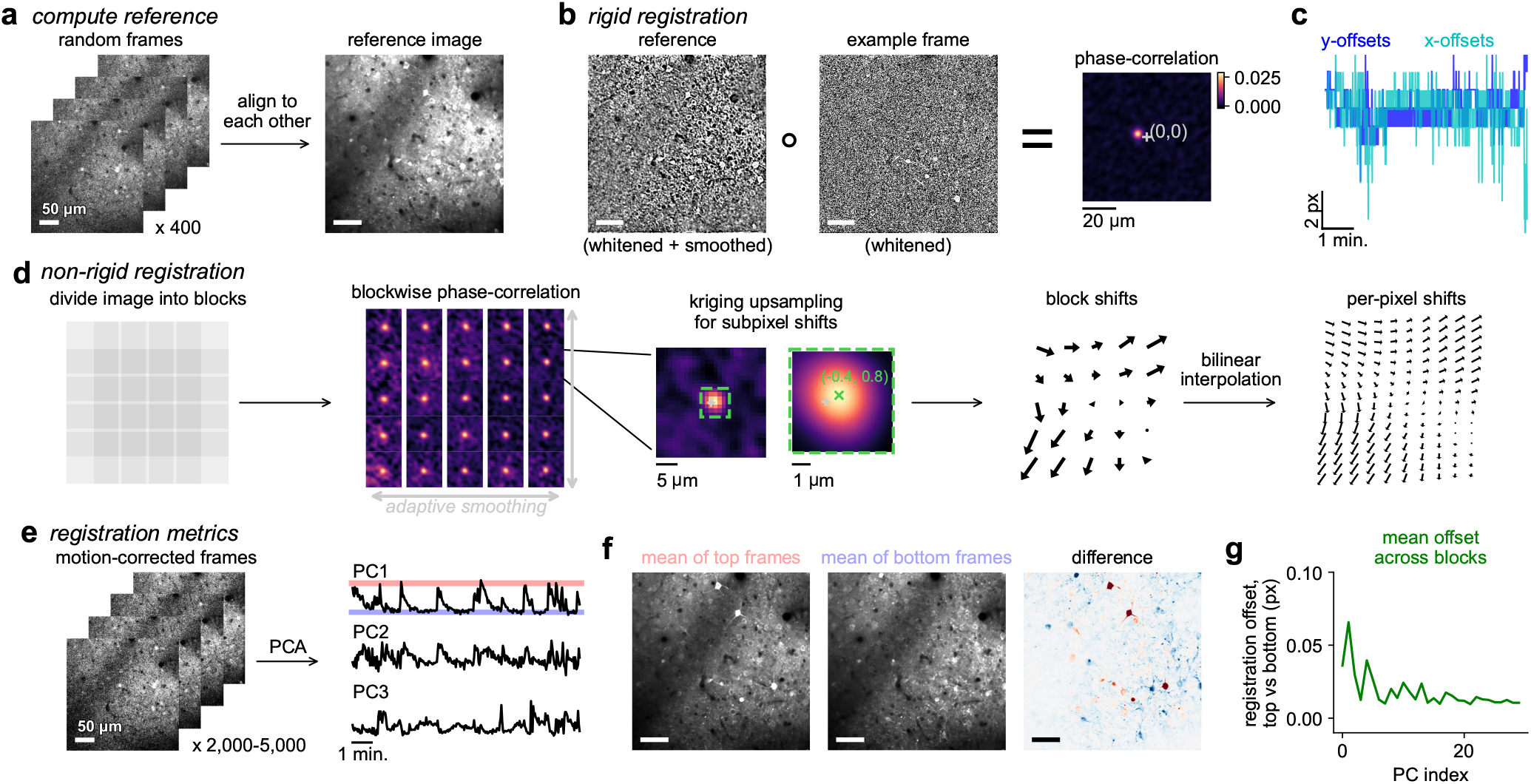
Motion correction and stability metrics. **a**, Reference image is computed by iterative re-alignment of a random subset of frames. **b**, Rigid motion offsets are computed by correlation of whitened reference and frame, i.e. phase correlation. **c**, Calculated X/Y rigid motion offsets. **d**, Non-rigid motion offsets computed by dividing image into overlapping blocks, computing phase correlation in each block, smoothing across blocks, taking the optimal shifts per block and upsampling them via bilinear interpolation. **e**, Registration performance is assessed by applied principal components analysis (PCA) to the motion-corrected frames. The PCs are expected to correspond to neural activity if all motion is corrected. **f**, Mean of the top PC1 frames (left), bottom PC1 frames (middle) and their difference (right). **g**, Registration offset lengths between top and bottom frame means.

**Figure 3:**
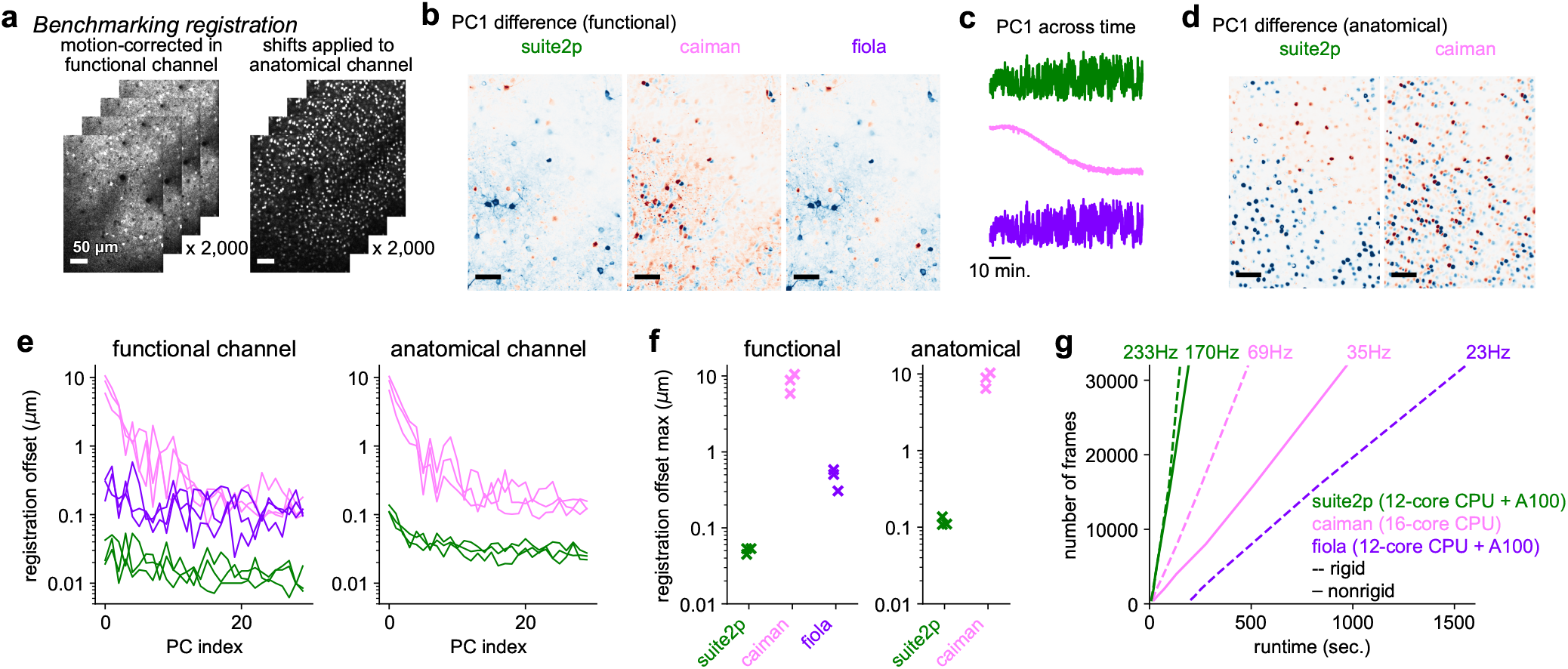
Benchmarking motion correction in vivo. **a**, Recordings of neurons expressing both GCaMP6s and tdTomato. Motion correction shifts were computed using the functional channel, and applied to the anatomical channel. **b**, Difference between top frames mean and bottom frames mean for PC1 (as in Figure 2f) for motion correction performed using Suite2p, Caiman, and Fiola, for the functional channel. **c**, Time course of PC1 across frames. **d**, Same as **b** on the anatomical channel. **e**, Registration offset lengths (like Figure 2g) for the functional (left) and anatomical channels (right). **f**, Maximum registration offset across the top 30 PCs for the functional (left) and anatomical (right) channel. **g**, Runtime performance. Frames per second reported as slope of linear fit.

**Figure 4:**
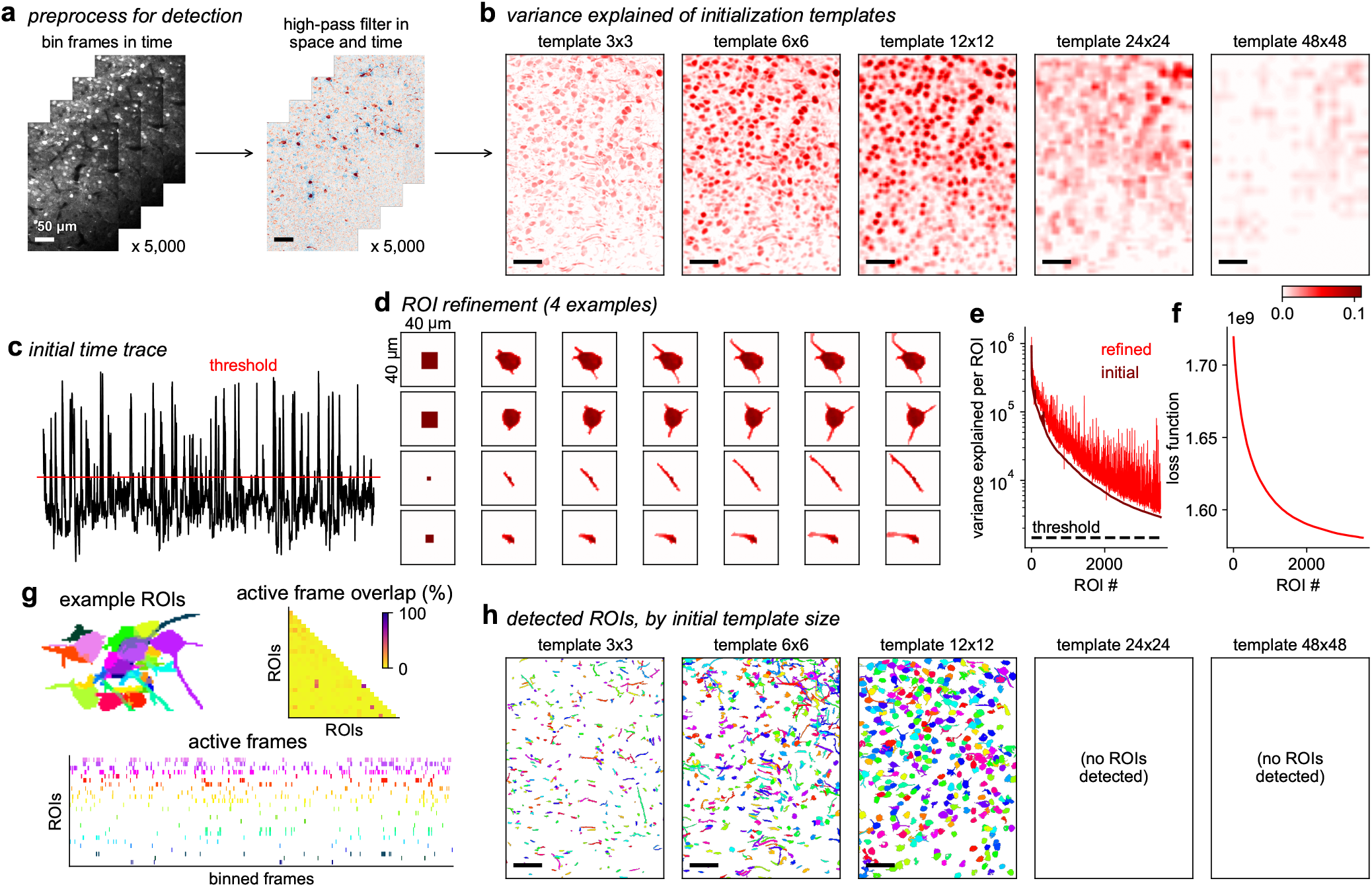
ROI detection. **a**, Frames are binned in time (left) and then high-pass filtered in space and time (right). **b**, Time-averaged thresholded activity from convolved square templates of different sizes. **c**, Initial time trace corresponding to the dot product of the template in **c** with each movie frame. **d**, Iterative template refinement for four example ROIs starting with the best fitting square templates from **b. e**, Variance explained per neuron shown both for the initialization template and for the refined template. **f**, Loss function as a function of the number of greedily extracted cells. **g**, Example region with many cells detected and their spatial overlaps (top left), temporal overlaps (top right) and active frames crossing the threshold (bottom). **h**, Detected ROIs divided by the sizes of their initial square templates.

### Motion correction

Brain motion on the order of several microns is very common in awake, in vivo recordings and thus it must be corrected computationally, ideally with sub-micron precision. Since the two-photon, pixel-by-pixel acquisition is sequential, brain position can change over the course of a single frame, resulting in a non-rigid deformation of the final frame [29]. Non-rigid deformations can also be induced by other factors, generally on longer timescales, such as very slight angle changes, laser beam walk and blood flow changes. These non-rigid deformations have to be addressed computationally in order to maintain sub-micron registration accuracy across the entire field of view.

Across biological applications, a large array of registration methods exist. To choose between them, we consider a number of factors unique to neural recordings: 1) low SNR / high shot noise on a frame-by-frame basis; 2) large number of frames per recording (hundreds of thousands); 3) non-rigid, but locally-smooth deformations. The first factor implies that we should take advantage of global motion estimation as much as possible, to integrate over the SNR of the entire frame. The second factor implies that processing should be fast, and the final factor implies that some amount of non-rigid correction must be done, even though locally there might not be sufficient information to estimate the motion due to the first factor.

To begin motion correction, it is necessary to create a reference image. The quality of the final registration depends critically on the resolution and SNR of this image. Due to low SNR per frame, a single frame cannot be used as reference. While an average of the first few hundred frames can be used [30–32], this average generally produces a blurry image due to the existence of brain motion (Figure S3a). To fix this problem, we refine the reference image iteratively from a subset of frames, averaging only frames that are well-aligned to the target at every iteration (Figure S3b, Figure 2a).

The second step computes a global correction for each frame, which we do using a rigid phase-correlation approach. Frame correlation approaches in general have the advantage they can be efficiently computed in the Fourier domain, and they integrate information across the frame to produce a single global registration offset. While previous methods like NoRMCorre and Caiman use a “cross”-correlation approach [30, 31], we found “phase”-correlation to perform substantially better ([33, 34], Figure 2b). The difference between these two is a whitening of the frequency spectrum in phase correlation, which emphasizes edge-like features in the image, rather than the low frequencies which are typically less informative about precise position.

After global, rigid correction per frame, the third step applies a non-rigid correction. This step compensates for the differential movements that different portions of the frame may contain, and thus is a relatively local, smaller correction. We determine this correction by dividing the frame into blocks, similar to other methods [30], and computing the phase correlation for each block. At this stage, we could directly correct each block, similar to previous methods [30]. However, such a correction may be quite noisy, because individual blocks contain a fraction of the global information available in the image, and optical recordings are generally made with large shot noise. Thus, we apply an adaptive smoothing step, which smooths the phase correlation maps across adjacent blocks, with a smoothing constant that depends on the estimated SNR for each block. Thus, blocks with high SNR will have an accurate and very local phase correlation map, while blocks with low SNR will have a more global map, but still be relatively accurate. Spatial offsets for each block are estimated from the peaks of the smoothed phase correlation maps after it has been spatially upsampled by a factor of 10 using Kriging interpolation with a 2D Gaussian kernel [35]. This upsampling enables subpixel corrections. The spatial offsets at each block are then interpolated to each pixel in the image, and applied to the frame using bilinear interpolation to generate the final corrected image.

### Registration benchmarks and quality control

To determine the success of motion correction, we designed a set of registration metrics that can be applied to any recording and any algorithm. Assuming all motion in the recording has been removed, the top principal components (PCs) of the corrected movie should be related to the activity of the somas or other compartments. If residual motion is left, it will correlate pixels over large spatial extents, and thus the motion will manifest in the top PCs (Figure 2e). We thus investigate the top PCs individually, by averaging the “top” frames with the largest projections onto the PC, and the “bottom” frames with the lowest projections (Figure 2f), and register these averages to each other. The amplitude of the offsets can provide an estimate of residual motion related to this PC (Figure 2g). We can also determine whether the residual motion is rigid or non-rigid, simply by estimating offsets with rigid/non-rigid registration. The difference between top and bottom frames can be inspected for qualitative assessment, and may provide insight into the type of residual motion present [36]. We make these metrics and corresponding visualizations easily available in the Suite2p GUI (Figure S1).

Beyond helping users understand the motion in their recordings, the registration metrics also enable a straightforward comparison between different algorithms on real data. We benchmarked Suite2p against Caiman [31], a popular calcium imaging processing tool, and Fiola [32], a more recent processing tool with GPU-accelerated registration. We ran Suite2p, Caiman and Fiola on a multi-ROI recording with both a functional and an anatomical channel (Figure 3a). We used the functional channel to determine motion offsets that were applied to both the functional and anatomical data and then computed the registration metrics (Figure 3b-d). Using the non-rigid registration offsets, we found that Suite2p outperformed Caiman and Fiola on the functional and anatomical channel metrics (Figure 3ef, Fiola cannot apply the anatomical channel correction and does not perform non-rigid motion correction). The metrics also indicated a gradual temporal misalignment for Caiman, which we verified by inspecting the offsets (Figure S4) and which we attribute to the incremental, “online” updating of the template image.

The Suite2p rigid and non-rigid registration algorithms were substantially faster than Fiola and Caiman, even when Caiman was run on a recent compute cluster node with 16 cores, and with Fiola running on the same GPUs as Suite2p (Figure 3g). We note that Fiola does not support non-rigid registration, and the speedups previously reported in relation to Caiman may have been due to the specific configuration and hardware used there [32].

### ROI detection

Once the movie is registered, the next step is ROI detection. Typical matrix decomposition approaches tend to extract many ROIs that are not single cellular compartments, such as neuropil and motion artifacts, because these sources induce correlated factors across pixels. While such factors can be classified post-hoc and eliminated, they can affect the detection and activity extraction of overlapping cellular compartments. In our experience, we find such artifactual ROIs to be easily identified by their lack of activity transients. Conversely, if an ROI has even a single high-activity transient, it likely corresponds to a cellular compartment. This led us to develop an algorithm that would specifically detect ROIs by their transient excursions above baseline. The algorithm can be formulated as matrix decomposition with dual L0 penalties on spatial and temporal factors (see Methods). However, it can also be described in more intuitive terms, which we do below.

To start, the registered movie is preprocessed by binning and high-pass filtering (Figure 4a). We then convolve each binned frame with square templates of different sizes and threshold at a fixed high value to identify transients, which are then summed up to generate a score for each template size, which is derived from the variance explained of a custom loss function (Figure 4b and Methods). The score corresponds to the data variance accounted for by the model if we were to introduce a square template of that size at that particular location. From here, the algorithm proceeds iteratively, identifying the highest peak across variance explained maps, and optimizing a template centered on that peak in order to explain more variance (Figure 4cd) which leads to a refinement of its shape. The activity of the ROI is then subtracted, the variance explained maps get updated efficiently, and the next ROI is extracted in a similar fashion, until no ROIs are left above a certain variance explained threshold (Figure 4ef), similar to matching pursuit algorithms [38]. The final reconstruction of example frames is shown in Figure S5.

This algorithm allows for overlapping ROIs to be found (Figure 4g). This is possible in cases of low to medium overlap, because individual neurons are typically sparsely active, resulting in a low number of collisions of simultaneous activity in pairs of overlapping templates (Figure 4g). This in turn allow the algorithm to find the true mask shape based on periods of high activity of individual ROIs. Additionally, the algorithm can detect various types of compartments like somas and dendrites (Figure 4h) due to its multi-scale nature.

To evaluate the algorithm, we generated hybrid ground-truth recordings based on real in vivo recordings. Our methodology relies on a riboL1-based calcium indicator which localizes to the soma, and thus makes cellular identification straightforward, even using anatomical methods like Cellpose (Figure 5a) [37, 39]. Similarly, traces were extracted just by averaging pixels (Figure 5b). To make the benchmark challenging, we added simulated neuropil, both as individual localized compartments (dendrites/axons) and as a spatially-diffuse, averaged signal (Figure 5a). The timecourses of these added components were obtained from real activity traces recorded in the other imaging planes. Finally, while the original recording was acquired at high laser powers, we added shot noise to make the hybrid ground-truth more representative of standard recordings.

**Figure 5:**
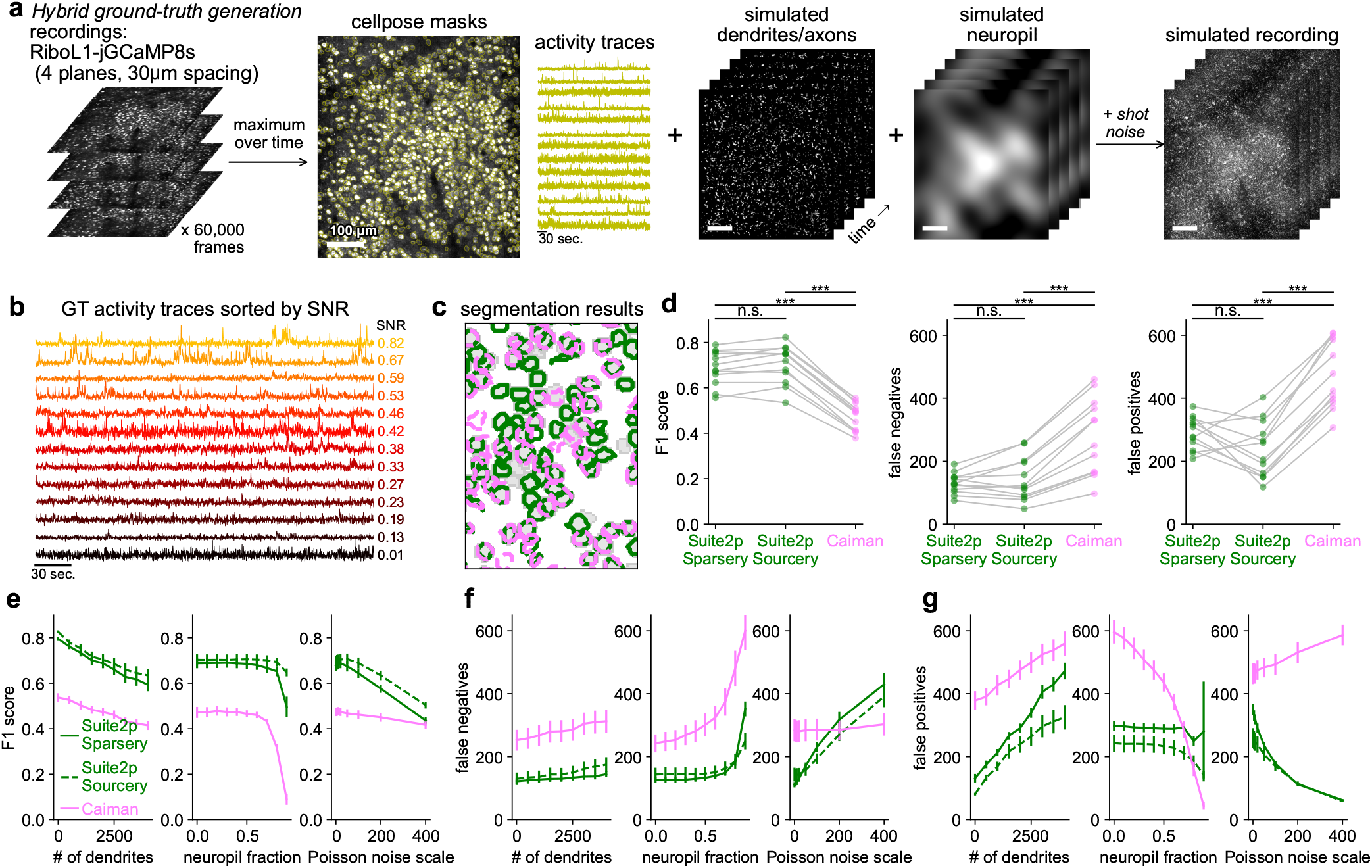
ROI detection benchmarks with hybrid ground-truth. **a**, Hybrid ground-truth generation using recordings of visual cortical neurons expressing riboL1-GCaMP8s [37]. Anatomically-detected ROIs were used as ground truth. We added simulated processes and neuropil using activity from other recorded planes. **b**, Example ground-truth activity traces from anatomically-detected ROIs sorted by SNR. **c**, Segmentation results from Suite2p (green) and Caiman (pink) overlaid on ground-truth ROIs (gray). **d**, F1 scores (left), false negatives (middle), and false positives (right) relative to the anatomical ground truth (paired two-sided Wilcoxon signed-rank test, n=12 simulations, *p*=4.9 ×10^™4^, *p*=4.9 ×10^™4^, and *p*=9.3 ×10^™3^. **e**, F1 scores across sweeps of simulation parameters. The values of 2,000, 0.4 and 20 respectively were used in **bc**. Error bars represent s.e.m, n=12 planes from 3 mice. **fg**, Same as **e** for false negatives and false positives.

We compared Suite2p to the CNMF-based [17] algorithm from Caiman, which was kept unmodified in Fiola for ROI detection (Figure 5c). Across recordings, Suite2p maintained a higher F1 score, with fewer false negatives and fewer false positives (Figure 5d).

The higher F1 score of Suite2p was maintained when varying the parameters of the simulation (Figure 5e-g). Suite2p also ran faster than Caiman, averaging 125 seconds for detection and extraction on the simulations, compared to 1104 seconds for Caiman. We note that we optimized a single parameter for both Suite2p and Caiman to finetune the total number of extracted ROIs, which had a small effect on performance (Figure S6). We also optimized together five Caiman parameters, which led only to very modest increases in performance (Figure S7), so we did not include this optimization in the main results to prevent overfitting.

Within Suite2p, we also include two other detection algorithms, Sourcery and Cellpose, which are useful in low-SNR scenarios and for different sensors (see Methods). We show the performance of Sourcery in the benchmarks (Figure 5d-g and Figure S6). We further evaluated Suite2p on the ground-truth datasets from [31]. The comparison is somewhat flawed,because these datasets were used as training data for Caiman. Furthermore, the ground truth was obtained by manual annotation, which is imprecise and may miss sparsely-active or low SNR cells. Nonetheless, the Suite2p algorithms performed well on this benchmark, in fact outperforming Caiman which was trained on this data (Figure S8). Finally, we show that the Suite2p algorithms can be applied on other types of data, such as interneurons recorded by calcium imaging (Figure S9ab), neurons recorded by calcium imaging through a one-photon miniscope in a freely moving rat (Figure S9c-e, compared to Caiman [31]) [40], and neurons recorded by voltage imaging in zebrafish (Figure S9fg, compared to Volpy [41]) [42].

### Extraction, deconvolution, GUI and cell tracking

The final steps in the pipeline are activity extraction, neuropil subtraction, and spike deconvolution. For the former, we extract ROI traces simply as averages of pixels, taking care to remove portions of each ROI that overlap with others as a conservative approach. In two-photon imaging, the point spread function (PSF) is larger in Z than in XY, and thus the ROI traces contain out-of-focus fluorescence from above and below the ROI. We estimate this out-of-focus fluorescence, i.e. “neuropil”, as the average activity of non-ROI pixels surrounding each ROI. This computation is independent of the ROI detection algorithm, and does not depend on the neuropil model used there. We subtract the neuropil signal from each ROI signal with a weighting coefficient of 0.7 for all ROIs, because estimating this coefficient per ROI can be noisy (Figure S10) [43]. Other methods like Caiman and Fiola use overlapping pixels and source deconvolution to separate ROI signals and background/neuropil signals during detection, which may result in biases (Figure S10). Similarly for spike deconvolution we adopt a simple algorithm, OASIS with the AR-1 model and no sparsity constraints [12,44].

Finally, the Suite2p GUI was designed to efficiently display multiple image projections, ROI metrics, correlations between ROIs, and population activity visualization via Rastermap [26] (Figure S1). The GUI can also be used to manually classify ROIs, but we recommend using this functionality sparingly, and focusing on changing algorithm parameters or recording configurations to obtain better automated results. In our experience, checking each recording in Suite2p immediately after acquisition is an invaluable step that allows for early detection and debugging of problems with the recording or the data processing.

For cell tracking over days, a common approach is to design algorithms that take the output of a pipeline such as Suite2p and Caiman, and then match ROI masks detected in different sessions as belonging to the same neuron according to various overlap criteria [45]. In such a two-step approach, the accurate tracking of neurons depends on the ROI extraction algorithm first finding the same cell across multiple sessions. Instead, we here demonstrate a “single-step” approach that works as well as or better than the two-step method. Our approach simply concatenates all data and runs Suite2p on the concatenated movie. This approach relies on the state-of-the-art registration algorithm in Suite2p (described above) to correct for any rigid or non-rigid movements across days. In our data, this approach was able to find a large number of ROIs in an example recording from hippocampal area CA1 while a mouse navigates a virtual reality environment (Figure 6a). The detected neurons had activity across days (Figure 6b). The single-step Suite2p approach tracked more neurons over days than a two-step pipeline called Track2p that uses the single-session Suite2p results (Figure 6c, 10,997 neurons tracked vs 5,836). When using the neuron-to-neuron correlation matrix as a benchmark of stability as in [45], we find that Suite2p matches the level of stability of Track2p (Figure 6de). As another benchmark of stability, we computed corridor tuning curves for each neuron, and evaluated their correlation across days, again finding similar stability as Track2p (Figure 6fg). Thus, the one-step tracking approach extracts neurons with similar stability as a dedicated two-step algorithm, while tracking substantially more ROIs (Figure 6c). We expect this approach to work well as long as the imaging plane is relatively well-aligned day after day. Good alignment can be obtained with the online motion correction algorithms which we provided in scanimage [46]. When the alignment between days degrades, we expect all algorithms to struggle with cell tracking, mainly because the same neurons are no longer in the image.

**Figure 6:**
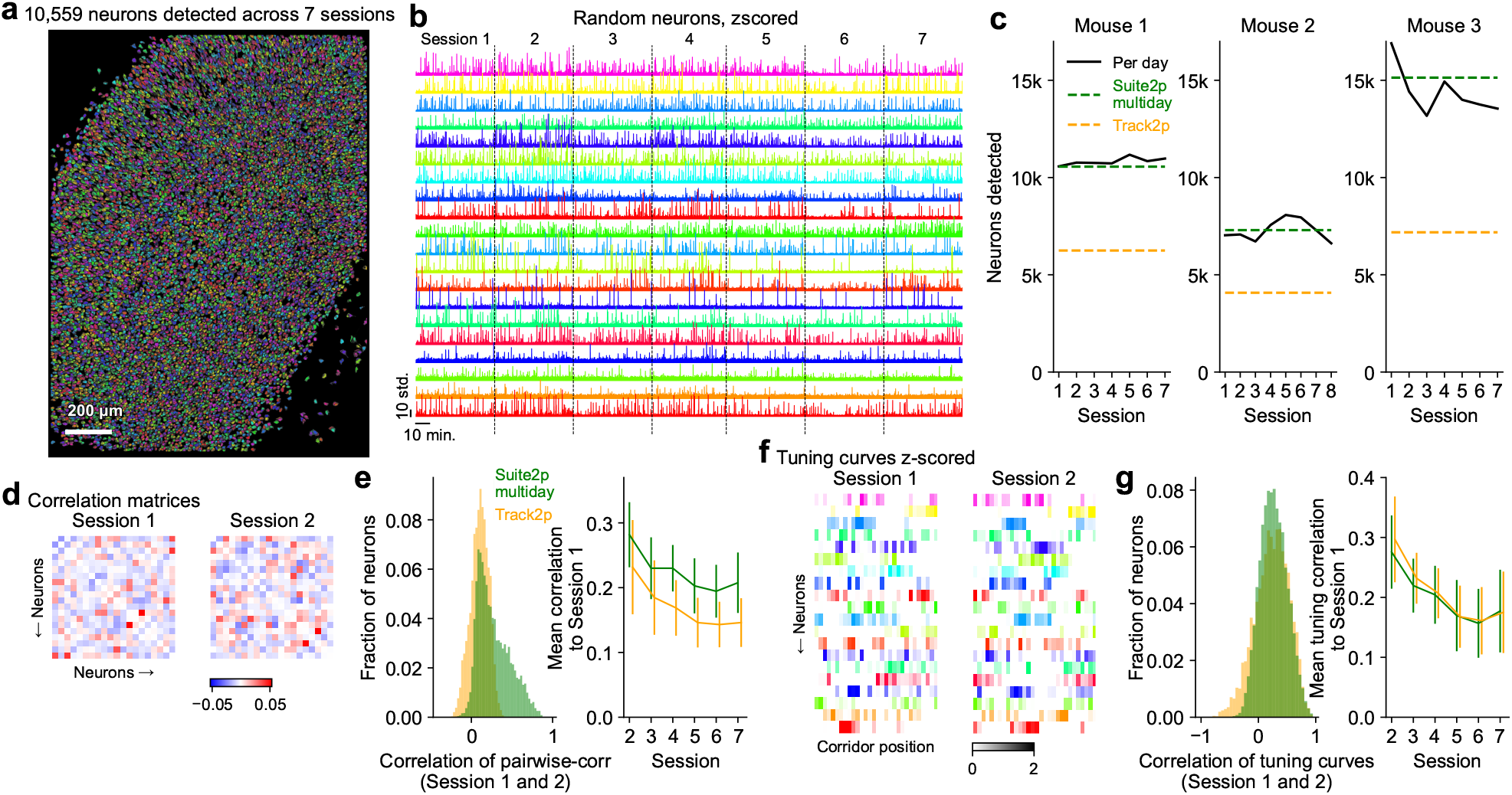
Cell tracking over days. **a**, Masks of neurons detected by Suite2p in concatenated recordings from 7 different days. **b**, Activity of cells whose masks are shown in **a. c**, Number of neurons tracked across days by the multiday Suite2p method vs a dedicated cell tracking approach (Track2p). **d**, Pairwise correlation matrices between neurons for two sessions (example submatrix). **e**, (left) Histogram of correlation values between rows of the pairwise correlation matrices between two example sessions. (right) Mean of these correlation values between session 1 and all other sessions, error bars represent s.e.m., n=3 mice. **f**, Z-scored single-corridor tuning curves for neurons in the first two sessions illustrated as heatmaps. **g**, (left) Histogram of correlation values between tuning curves computed from two example sessions. (right) Mean of these correlation values between session 1 and all other sessions, error bars represent s.e.m., n=3 mice.

## Discussion

Suite2p provides a complete suite of algorithms for extracting neural activity from optical recordings. As a practicing lab, we perform several large-scale neural recordings every day, which has allowed us to thoroughly test the robustness of the pipeline. Using the Janelia GPU cluster and parallelizing the data processing over planes, we now obtain high-quality processed datasets within 20 minutes of a recording finishing. This fast feedback allows for quick experimental iterations and faster debugging and improvement of recording configurations.

The fast registration algorithms in Suite2p are also integrated with Scanimage [46] to provide realtime Z-drift estimation and correction. We have found this step to be crucial in maintaining stability over long recording durations. Suite2p also aligns dense z-stacks if volumetric data is provided, and the results can be visualized in the GUI.

## Acknowledgments

This research was funded by the Howard Hughes Medical Institute at the Janelia Research Campus. From the Vivarium, we thank Jim Cox, Crystall Lopez, Anne Kuzspit, Miriam Rose, Alexa Gracias, Gillian Harris, Sarah Lindo, and their respective teams for animal breeding, husbandry, and surgeries. From JeT, we thank Daniel Flickinger, Vasily Goncharov, Alex Sohn, Tobias Goulet, and Steven Sawtelle for help with rig maintenance and upgrades. We thank Sverre Grødem, Kristian K Lensjø, and Marianne Fyhn for the RiboL1-jGCaMP8s virus. We thank the GENIE project team, Caiying Guo and Xianling Zhao at Janelia for generating the TetO-jGCaMP8s transgenic mice. From MBF Bioscience we thank Georg Jaindl, Mitchell Sandoe, and Boris Djiguemde for scanimage support. We thank members of our lab for testing the latest Suite2p version. We thank Kenneth Harris and Matteo Carandini for supporting the development of an earlier version of Suite2p, and Sylvia Schroeder and Mario Dipoppa for testing and contributions to an earlier version. For testing an earlier version, we also thank Federico Rossi, Ross Williamson, Dimitar Kostadinov, Henry Dalgleish, Michael Hausser, Alice O’Leary and Asaph Zylbertal. We thank Minh Thao Nguyen, Andrea Pierré, Joao Couto, and Tolga Dincer for code contributions to improve file format support. We thank Jan Willem de Gee for contributing an option for running motion correction multiple times, and Seetha Krishnan for contributing a manual ROI drawing tool. We thank the authors of [31, 40, 42] for sharing their datasets.

## Data availability

The data is available at janelia.figshare.com/ articles/dataset/Dataset_for_Extracting_ large-scale_neural_activity_with_Suite2p_/31460065 [47].

The one-photon and voltage imaging recording are publicly available on zenodo [40, 42, 48, 49]. The Caiman ground-truth data is also publicly available on zenodo [31, 50].

## Code availability

The code and GUI for Suite2p are available at https://www.github.com/mouseland/suite2p. Scripts for recreating the analyses in the figures are available at https://github.com/MouseLand/suite2p/tree/main/paper/.

## Methods

The Suite2p code library is implemented in Python 3 [51], using numpy, scipy, pytorch, numba, scikit-learn, tifffile, and ScanImageTiffReader for processing [52– 58], and pyqt, pyqtgraph, and superqt for the GUI [59–61]. The figures were made using matplotlib and jupyter-notebook [62, 63]. All experimental procedures were conducted according to IACUC, ethics approval received from the IACUC board at HHMI Janelia Research Campus.

### Data acquisition

#### Animals

All experimental procedures were conducted according to IACUC. We performed recordings in one C57Bl/6Crl mouse, three VGAT-CRE x Ai14 mice (JAX 016962, JAX 007914), one mouse bred to express jGCaMP8s in excitatory neurons: TIGRE2-RiboL1-jGCaMP8s-IRES-tTA2 (variant of JAX039267 without IRES and with WPRE) x Slc17a7-cre (JAX 023527) [28], and three mice bred to express GCamP6f in excitatory neurons: Thy1-GCaMP6f GP5.17 mice (JAX 025393) [64]. These mice were male and female, and ranged from 2 to 12 months of age. Mice were housed in reverse light cycle, and were housed with siblings before and after surgery. Holding rooms are set to a temperature of 70°F +/-2°F, and humidity of 50%rH +/-20%.

#### Surgical procedures

Surgeries were performed in adult mice (P56–P200) following procedures outlined in [65]. In brief, mice were anesthetized with Isoflurane while a craniotomy was performed. Marcaine (no more than 8 mg/kg) was injected subcutaneously beneath the incision area, and warmed fluids + 5% dextrose and Buprenorphine 0.1 mg/kg (systemic analgesic) were administered subcutaneously along with Dexamethasone 2 mg/kg via intramuscular route. Measurements were taken to determine bregma-lambda distance and location of a 4 mm circular window over visual cortex, as far lateral and caudal as possible without compromising the stability of the implant. In VGAT-CRE mice, injections were performed of soma-targeted RiboL1-jGCaMP8s (a mixture of AAV9-Thy1s:TTA and AAV9-TRE3G:RiboL1-jGCaMP8s) [37, 66]. In the C57 mouse, injections were performed of AAV-hSyn1-GCaMP6s-P2A-nls-dTomato, (Addgene plasmid # 51084 ; http://n2t.net/addgene:51084;RRID:Addgene_51084). A 4+5 mm double window was placed into the craniotomy so that the 4mm window replaced the previously removed bone piece and the 5mm window lay over the edge of the bone. In the Thy1-GCaMP6f mice, we placed a window over hippocampal CA1. For this, the craniotomy was centered at 1.8 mm AP and 2.0 mm ML from bregma, then cortex was aspirated, and a 3 mm glass coverslip attached to a stainless-steel was implanted over the dorsal CA1 region, similar to [67]. After surgery, Ketoprofen 5mg/kg was administered subcutaneously and the animals allowed to recover on heat. The mice were monitored for pain or distress and Ketoprofen 5mg/kg was administered for 2 additional days following surgery.

#### Imaging acquisition

During the recordings, mice were free to run on an air-floating ball or a treadmill. Mice were acclimatized to running head-fixed for several sessions before imaging.

We used ScanImage [46] for data acquisition, and a custom online Z-correction module to correct for Z and XY drift online during the recording, called the “MariusMotionEstimator” and the “MariusMotionCorrector” in ScanImage, this module was used for all recordings except the recording in Figure 2 for benchmarking motion correction.

For the many plane recording (Figure 1), we used a commercial two-photon ThorLabs Bergamo2 microscope with a piezo for multiplane acquisition, an acquisition system from MBF Bioscience (the “slow” vDAQ), and an objective with lower zoom and NA (Nikon CFI Plan Apo 10XC Glyc, 0.5 NA, WD 5.5mm, MRD71120). We used a Toptica Femtofiber 920nm laser customized to a lower repetition rate of 30Mhz and shorter pulse width (100fs) with GDD compensation, with a max power at the sample of approximately 240mW, which improved recording quality. We recorded eight planes one of which was a flyback plane, at a rate of 1.87Hz, with pixel sizes of 2 *µ*m in both X and Y, with the automated XYZ drift correction turned on (Figure S2). The distance between the planes was 30 *µ*m, which reduced the likelihood of detecting cells on multiple planes. The laser pulses were synchronized with scanimage, allowing for precise matching of pulses to pixels. While possible on this setup, we did not use the uniform sampling mode. During the experiment, the mouse ran in a closed-loop virtual reality with six different corridors. When the mouse ran consistently, then the corridor along with the following gray screen was traversed in 12 sec., or approximately 22 imaging frames. We interpolated to this length for computing single-neuron tuning curves to the corridors.

For the benchmarking experiments, we used a custom-built 2-photon mesoscope [68] to record neural activity. For the registration benchmarking, the XYZ drift correction was turned off. This recording had a pixel resolution of 1 *µ*m, three fields of view of size 1112 x 650 pixels, a recording rate of 6.76 Hz, and a recording length of 80 minutes. We also used this recording for Figure 1b, Figure 2, Figure 3, Figure 4, Figure S10.

For the detection benchmarking experiments, we used an upgrade of the mesoscope that allowed us to approximately double the number of recorded neurons using temporal multiplexing [69], as described in [65], with XYZ drift correction turned on. The recordings were at a pixel resolution of 1 *µ*m, with four planes spaced 30 *µ*m apart, a frame rate of 15-15.5Hz, a frame size that varied slightly across the three mice due to manual selection of the recording field of view (744×654, 726×654, and 748×652), and a recording length of 60-66 minutes.

For the interneuron experiment, we used the mesoscope to perform a recording in a VGAT-CRE mouse at 28Hz frame rate with multiplexing, with a pixel resolution of 2 *µ*m. First, both the green and red channels were acquired with two PMTs for 1.5 minutes. Then only the green PMT was used for the functional recording for 35 minutes (this reduces the data storage required for such recordings).

For the tracking experiments, we used the ThorLabs Bergamo2 microscope with the Nikon 10X objective at 20Hz frame rate (mice 1 and 2), and the custom mesoscope with multiplexing at 10Hz frame rate (mouse 3), with a pixel resolution of 1.33 x 2 *µ*m. Neural activity was recorded across days while the mice performed a two-choice task in a closed-loop virtual reality with two corridors in which one corridor delivered rewards (similar to [70]).

### Overall pipeline settings

There are several settings, saved in ‘settings.npy’, that the user is recommended to modify for their experiment:

- `torch_device`: Torch device for performing operations, GPU is ‘cuda’ and CPU is ‘cpu’
- `tau`: Exponential timescale of decay of indicator in seconds, which is used in spike deconvolution algorithm in OASIS [12]. We recommend a timescale of 0.1/0.15/0.25 sec. for jGCaMP8 f/m/s and 0.5/0.75/1 sec. for GCaMP6 f/m/s, based on benchmarking analyses similar to [44]. However, as shown there, the exact value of this parameter does not matter much.
- `fs`: Sampling frequency of each plane in Hz, i.e. the volume rate of the acquisition.
- `diameter`: Expected ROI diameter, used for detection with Cellpose or original Suite2p algorithm, Sourcery.

#### Input formats and settings

Suite2p supports tiffs, hdf5 files, NWB files, bruker tiffs, avi files, and dcimg files. Metadata about the recording and file saving locations is provided in a dictionary ‘db.npy’, or through the GUI, which is then converted to an npy file. These settings are used to convert the inputs into binary files which are memmapped during the processing. Each imaging plane (and Scanimage ROI - if performing multi-ROI acquisition) is saved as a separate binary file in a separate folder ‘planeX’, where X is a number iterated over ROIs and planes. The input settings include

- `data_path`: List of folders with tiffs or other files to
- `input_format`: Can be [‘tif’, ‘h5’, ‘nwb’, ‘bruker’, ‘movie’, ‘dcimg’].
- `keep_movie_raw`: Whether to keep binary file of non-registered frames.
- `nplanes`: Each tiff / file has this many planes in sequence.
- `nrois`: Each tiff / file has this many different ROIs.
- `nchannels`: Specify one- or two-channel recording.
- `functional_chan`: The channel to use to extract functional ROIs (1-based).
- `lines`, `dy`, `dx`: Line numbers for each ScanImage ROI within the tiffs, Y position for each ROI, and X position for each ROI.
- `save_path0`: Directory to store results, defaults to data_path[0].
- `fast_disk`: Directory to store temporary binary file (recommended to be SSD), defaults to save_path0.
- After conversion to binary files, a plane-specific ‘db.npy’ file is added to each plane folder with plane-specific keys, including:
- `Ly`, `Lx`: Size of the field of view in pixels in Y and X.
- `reg_file`: Location of the binary file, in which registered frames will be saved (before registration, this contains the raw frames).
- `raw_file` (optional): If `keep_movie_raw`, the location of the raw binary file to be registered and saved as `reg_file`.

### Motion correction

The settings for motion correction are in the dictionary called ‘registration’. The `batch_size` parameter sets the number of frames in each chunk processed together - this may need to be reduced on GPUs with less memory. There is the option to use the anatomical channel for registration, by setting `align_by_chan2` to True. During motion correction, all registered frames are cumulatively averaged, producing a registered mean image, saved in the registration outputs as `meanImg`, and `meanImg_chan2` if there is a second anatomical channel.

#### Bidirectional phase offset (optional)

If `do_bidiphase` is True, then the bidirectional phase offset is estimated from a subset of frames, with the size of the subset defined by the parameter `nimg_init`. The bidirectional phase offset is defined as the optimal shift between subsequent lines of the raster-scanning, computed via phase-correlation. Alternatively, the user can specify the shift with the setting `bidiphase`. The computed or user-defined shift is then applied before any motion estimates from frames, and when the motion correction shifts are applied to each frame. The shift used is saved in the registration outputs as key `bidiphase`.

#### Reference image computation

A subset of frames are read in from the entire movie at equal spacing, with the size of the subset defined by the parameter `nimg_init`. The pairwise correlations between these frames is computed (Figure S3b). We choose the frame which has the largest correlation with its 20 most-correlated frames, and average these 20 frames. We use this average as an initial reference image, and compute the phase-correlation (rigid) of all the frames in the subset to this reference image iteratively, across 8 iterations. On each iteration, we average the frames most correlated to the current reference, after applying the estimated shifts from the phase-correlation, to create a new reference. The number of top correlated frames to be averaged is a function of the iteration *i* defined as nimg_init *(1 + *i*)*/*16. This reference image is saved in the registration outputs as key `refImg`.

#### Rigid registration

The rigid registration step computes the offset in X and Y between each frame and the reference image, and applies these shifts to each frame. To compute the offset, the reference image is first clipped, whitened and smoothed. The clipping, done if `norm_frames` is True (default is True), computes the first and 99th percentile of the pixel intensities in the reference image, and clips the reference image at these values, saved as keys `rmin` and `rmax` in the registration outputs. This clipping is also applied to all frames in the movie before computing the motion shifts. The reference image is next transferred to the Fourier domain and whitened. Filtering by a Gaussian of standard deviation `smooth_sigma` is done directly in the Fourier domain by elementwise multiplication of the Fourier transform of this Gaussian.

Next we apply a spatial tapering on each frame of the movie, so that the phase-correlation is focused on the central part of the field of view rather than the edges. We define the spatial taper as the outer product of two sigmoids **m**_*y*_ and **m**_*x*_ across *y* and *x* pixels. The slope of the sigmoid is controlled by setting `spatial_taper` *t*:

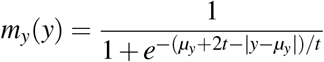

where pixel center *µ*_*y*_ defined as (*L*_*y*_ ™1)*/*2.

Then we whiten each frame and compute the cross-correlation of the reference with each whitened frame; cross-correlation using whitened images is called “phase-correlation” (Figure 2b). The fast FFT operation is greatly accelerated using the GPU.

The peak of the phase-correlation is determined in a range up to a fraction of the frame size, defined by `maxregshift`, to avoid outliers. The peak of the phase-correlation defines the offset in X and Y of the frame with the reference. We shift each frame according to the offsets using the ‘roll’ operation for efficiency - this fills the edges of the frame with the opposing edges during shifts, but we later ignore these regions during cell detection anyway.

#### Non-rigid registration

After each batch of frames is rigid registered, we optionally compute and apply non-rigid shifts to each frame, if `nonrigid` is True. We recommend applying this step on most recordings, especially if they have a sufficient SNR. One can tell if a recording has sufficient SNR by the outliers of the registration process: if the registration offset traces are jagged and jump to extreme values and back, this is due to low SNR (or possibly due to extreme instability in the recording setup). The non-rigid shifts are computed by performing phase-correlation on square sub-blocks of the frame defined to be overlapping by at least 50% in each dimension, with the size defined by the setting `block_size` (Figure 2d). Each block is spatially tapered with a sigmoidal function with slope parameter 2*`smooth_sigma` and whitened, and the phase-correlation is computed with the reference image in each block (filtered and whitened as in rigid registration). The maximal allowed shift of each block is defined in pixels by the setting `maxregshiftNR`.

Some parts of the image on some frames may have lower SNR - therefore we apply adaptive smoothing of the phase-correlation maps across blocks as a function of SNR. The smoothing filter is a 2D Gaussian with standard deviation equal to one block, that is applied up to two times. The SNR is estimated as the ratio between the peak value of the phase-correlation and the maximum phase-correlation outside the peak region (3×3 pixels). If the SNR is less than the `SNR_thresh` setting, then the smoothed phase-correlation is used. If the once-smoothed phase-correlation still does not have SNR higher than the `SNR_thresh`, then the twice-smoothed phase-correlation is used.

To estimate the XY shift of each block, we upsample the phase-correlation map by a factor of 10 around the peak (3×3 pixels) using Kriging interpolation with a Gaussian kernel of standard deviation 0.85 pixels. The peak of the upsampled correlation map provides a subpixel estimate of the XY shift for each block. To convert from block-wise shifts to a shift for each pixel in the image, we use bilinear interpolation, assuming the block shifts correspond to the shifts of their center pixels. We then shift the image by the pixel-wise shift map using bilinear interpolation. These interpolation operations are fast to perform on the GPU.

#### Valid region estimation

Because the frames have moved in X and Y, there are regions on the edges of the frames that are missing information as the putative ROIs have moved outside the field of view. We thus ignore these edge regions for ROI detection. We compute the edge regions using the offsets from the rigid registration, excluding frames with shifts which may be outliers (called `badframes` in the output of the registration, these frames are also excluded from ROI detection and can be user-defined). For this we median filter the XY-offsets and the peak phase-correlation in time with a window of 101 frames. We define large deviations using the Euclidean distance between the XY-offsets and the median-filtered XY-offsets, normalized by the mean distance. We define low correlation frames by the ratio between the peak phase-correlation and the median-filtered peak phase-correlation. If the ratio of these two values is larger than 100*`th_badframes`, then the frame is defined as a ‘bad’ frame and excluded from the valid region computation. We also define bad frames to include frames in which the shift was larger than 95% of the maximal allowed rigid shift, defined by `maxregshift`. Excluding the bad frames, the maximum absolute shifts in both X and Y are computed across frames. These pixel shifts define the sizes of the excluded edges on each side of the frame.

#### Registration metrics

We subsample 2,000-5,000 frames from the registered recording to estimate the accuracy of the motion correction (Figure 2e). We only sample 2,000 frames from the recording if the frame size is larger than 700 (to avoid memory issues) or if there are fewer than 5,000 frames. We crop the frames to the valid frame region and then compute the top 30 principal components of the frames. We average the frames with the 300 smallest weights and the 300 largest weights for each PC (Figure 2f). We register these top and bottom mean frames to each other to determine the motion shifts within the PC. We return the rigid shift distance, the non-rigid shift distance averaged across blocks, the maximum non-rigid shift distance across blocks, and the rigid and non-rigid shift distances added together and averaged across blocks (shown in Figure 2g, as the ‘mean registration offset’).

### ROI detection

The settings for detection are contained in the dictionary ‘detection’. First, using the valid region and excluding the bad frames, the frames of the recording are binned. The bin size is defined as the rounded timescale of the indicator decay, in frames (`fs` * `tau`), since the frames within this bin size are likely to be correlated and contain redundant information. We create a binned movie of size up to `nbins`. If the movie is longer than `nbins` * `fs` * `tau`, then the bin size is defined as the number of frames divided by `nbins`.

Optionally the binned movie can be PCA-denoised, if `denoise` is True. The PCA denoising is performed in blocks of size `block_size` (registration setting). The denoising is the reconstruction of each block using the top half of the principal components computed per block.

Next the movie is high-pass filtered in time, to reduce slow temporal components such as drift or neuropil, by subtracting the Gaussian filtered version of the movie from itself, with a Gaussian of standard deviation `highpass_time`. After high-pass filtering, the maximum of the movie across time is computed as the maximum projection image, called `max_proj` in the detection outputs.

We provide three algorithms, which can be specified by the `algorithm` setting, as ‘sparsery’, ‘sourcery’, or ‘cellpose’. For Sparsery and Sourcery, we compute the per-pixel standard deviation of the difference between consecutive binned frames in time, which is an estimate of the shot noise per pixel.

After ROI detection, there are optional steps that can remove ROIs before signal extraction. By default, we remove ROIs with more than 75% of their pixels overlapping with other ROIs (specified as `max_overlap` = 0.75). Optionally, the user can specify minimum and/or maximum sizes for removing ROIs, defined by `npix_norm_min` and `npix_norm_max` (see more details about ‘npix_norm’ below). Also, a shape- and size-based classifier can be applied to the ROIs before extraction, with the `preclassify` setting specifying the threshold for ROI inclusion.

#### Sparsery algorithm

The main detection algorithm, Sparsery, in Suite2p uses functional information to detect ROIs. The settings for Sparsery are in the ‘sparsery_settings’ dictionary. We first remove the low spatial frequency components from the movie, which are the background neuropil signals (Figure 4a). We divide each binned frame by the per-pixel shot noise estimate and then high-pass filter each binned frame by subtracting the frame uniform filtered with a filter size in pixels of `highpass_neuropil`. The Sparsery algorithm performs a matrix decomposition on this movie, assuming L0-sparse sources in space and L0-sparse traces in time. The cost function is:

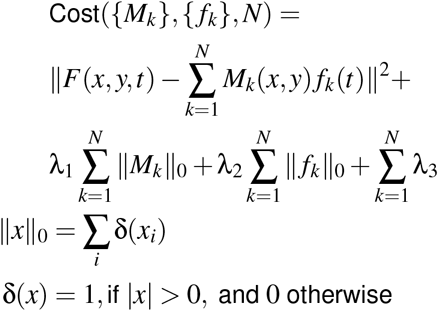

The effect of this cost function is to introduce a fixed penalty λ_1_ when activating a pixel in any ROI mask *M*_*k*_ and another fixed penalty λ_2_ when activating a temporal value in any ROI trace *f*_*k*_. Rather than directly specifying λ_1_ and λ_2_, we set them implicitly and adaptively in terms of the thresholds they induce on the activation of non-zero coefficients in the cost function. We optimize this function greedily, one ROI at a time. To initialize ROIs, we consider all possible uniform square templates of sizes 3×3, 6×6, 12×12, 24×24, 48×48. We found these values to cover most if not all sizes of ROIs in typical recordings. These serve as initial guesses for *M*_*k*_, and they explain a sufficient amount of variance to significantly reduce the cost function and initialize the optimization for *M*_*k*_. They also have the property that we can efficiently compute the variance explained of each square template via convolution with each frame, followed by thresholding according to λ_2_. For the larger templates, performing this convolution densely would incur large computational penalties. However, the variance explained of larger square templates is highly correlated across space: shifting a square template over by a few pixels from its optimal placement still explains a large amount of variance. Thus, we apply the square templates only at a subset of positions. We devised an efficient algorithm to do so via successive downsampling of the frames by additional factors of 2, followed by convolution with 3×3 uniform filters to exactly calculate the variance explained of templates spaced 2^*n*^ pixels apart where *n* is the downsampling iteration accounting for the square filter sizes 3 ·2^*n*^.

After constructing these variance explained maps at multiple resolutions, we estimate the scale on which most templates exist. The maximum of each template map across time is interpolated to the original movie size in pixels using bivariate spline approximation of degree 3 with the RectBivariateSpline function in scipy. Then the maximum across these interpolated maps is computed - is saved as `Vcorr` in the detection outputs and shown in the GUI as the ‘correlation map’, which can be used to determine if the algorithm may have missed correlated pixel groups. The peaks of this correlation map are found by subtracting off the maximum filtering of the correlation map with a filter size of 11×11 pixels and finding the pixels within 1e-4 of the maximum-filtering. The top 50 peaks, according to their value in the correlation map, are used to determine the best spatial scale, defined as the template map across scales on which the most of these 50 peaks have their maximum. Alternatively, the user can specify the estimated spatial scale (`spatial_scale`) with an integer from 1 to 4 corresponding to templates of size 6×6 to 48×48.

To compute the temporal threshold ‘Th2’ above which temporal coefficients are activated (corresponding to the λ_2_ penalty above), we multiply the inferred spatial scale by a constant: Th2 = 5 * `spatial_scale` * `threshold_scaling`, where the `threshold_scaling` can varied to find more or fewer ROIs. Intuitively, if the same cell is recorded at a higher pixel resolution, there will be more pixels per cells corresponding to a larger overall variance explainable for the same cell, hence the dependence on `threshold_scaling`. Only deviations above the Th2 threshold are used to compute the variance explained of the template (example trace in Figure 4c; variance explained maps in Figure 4b). To extract another cell, we require the square template to cross threshold at least a few times during the recording.

This is enforced by setting a minimum variance explained requirement of Th2· max(1, nframes*/*1200), where nframes is the number of binned frames. Thus,if the number of binned frames is relatively small, a single threshold crossing is necessary to initialize a template.

The algorithm iteratively detects ROIs to minimize the cost function. On each iteration, the location of highest variance explained across all template maps and positions is found and used as the initial template (Figure 4b). The active frames for the template are defined as the frames in the movie in which the template surpassed the threshold ‘Th2’ (Figure 4c). Alternatively, the user can specify a percentile to use instead of a threshold with parameter `active_percentile`, which can produce less sparse traces. The ROI is extended iteratively by one pixel on each edge, until there are no pixels that can be added (Figure 4d). Pixels are added if their mean intensities across active frames (called ‘lam’) are larger than the one-fifth of the maximum mean intensity across current ROI pixels; pixels are removed if they are less than one-fifth of the maximum. This procedure implicitly determines λ_1_, though in an adaptive manner for each ROI. The active frames are then recomputed using the newly refined ROI, and this process of iterative refinement is repeated twice more, or discontinued if there are no active frames after refinement. The activity of the ROI on its active frames are subtracted from the movie, and the variance explained of the templates are updated accordingly. The algorithm continues until the highest variance explained is below the threshold described in the previous paragraph, or when the maximum number of ROIs is reached, defined by parameter `max_ROIs`.

On each ROI iteration, there is a check to see if splitting the ROI would improve the variance explained. This check is performed by running iterative k-means, initialized with a split of the ROI into pixels above or below the mean pixel values at the time frame with the largest activation. Each ROI is stored as a dictionary, with the pixels of the ROI saved as ‘ypix’ and ‘xpix’ keys. The pixel weights ‘lam’ are the mean intensity of each pixel during the active frames, multiplied by the previously divided shot noise estimation. The initial template center location is saved in the ‘med’ key. The initial template map ID is saved in the ‘footprint’ key.

Note that this procedure is a very greedy, but very efficient optimization of the initial cost function described above. However, the algorithm maps onto intuitive steps that a human might take to find ROIs in such a movie: use a variance explained map to find regions of high unexplained variance, jump to frames where those regions are activated substantially, inspect whether the activated pixels at those frames look like cells, average the profile across frames, then account for the times when this cell is active and look for other masks that may be highly active at different times. Critical to these steps is the L0 interpretation: sources are only interpreted as cells or “good ROIs” if they are strongly active on at least a few frames, and highly spatially localized. Unfortunately, L0 optimization is very difficult, very slow, and always approximate, hence the need to make the compromises we described above.

#### Sourcery algorithm

Another detection algorithm, Sourcery, also uses functional information to detect ROIs. This algorithm can be useful for low SNR data with compact ROIs, as it relies less on sparse, transient deviations from a baseline and more on overall correlations between pixels. The cost function for Sourcery is similar to that above for Sparsery, excluding the temporal sparsity penalty:

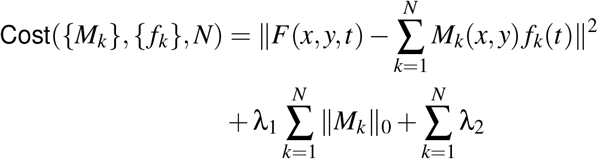

The optimization proceeds in a similar greedy fashion, however without consider temporal sparsity, and replacing some of the updating steps in Sparsery such as the splitting condition and the cell-by-cell update of the cost function. The settings for Sourcery are in the ‘sourcery_settings’ dictionary. The first step in Sourcery is to orthogonalize the binned movie via PCA (this movie has also been high-pass filtered in time as described above). We also divide each binned frame by the per-pixel shot noise estimate. Then the movie is smoothed spatially by a 2D Gaussian of standard deviation `diameter` / 10. The temporal PCs are computed by applying PCA to the covariance across timepoints, and projected onto the movie to obtain the spatial pixel-wise PCs (called *U* ). The number of top PCs used in the subsequent steps is the minimum between the number of time bins in the binned movie and one-half of the number of total pixels (X*Y) (usually the number of bins is smaller).

We model the neuropil as an orthonormal spatial basis functions (*S*) with low spatial frequency using a sine and cosine basis, and project out this basis from the spatial components of the data *U* . This is to improve cell detection and to help avoid extracting neuropil components with large spatial extents. The neuropil basis functions are defined as the outer products of sines and cosines, with the frequencies ranging from one to ⌊*n*_*y*_*/*2 ⌊and ⌊*n*_*x*_*/*2 ⌊cycles across the field of view, where *n*_*y*_ and *n*_*x*_ are the number of sines and cosines in Y and X respectively. *n*_*y*_ is set as the number of pixels in Y of the FOV divided by the `ratio_neuropil` setting and divided by the `diameter`; *n*_*x*_ is computed the same way using the number of pixels in X. The `ratio_neuropil` setting is approximately the spatial scale of the neuropil relative to the size of the ROIs - in many two-photon recordings this may be around 6. The basis functions are normalized to be unit norm. The neuropil contribution to the PCs (*N*) is estimated by linear regression, and subtracted from the PCs: *U*_cell_ = *U* ™*S*(*S*^⊤^*S*)^™1^(*S*^⊤^*U* ).

From this remaining *U*_cell_, the cell ROIs are found via an iterative algorithm. *U*_cell_ is first smoothed with a Gaussian filter with standard deviation of `diameter` / 4, producing *U*_smooth_. The sum of squared PC values for each pixel of the smoothed PCs is normalized by the sum of squared PC values from the non-smoothed PCs smoothed by the same Gaussian filter. A morphological opening is performed on this spatial map of radius `diameter` / 4 and subtracted off, in order to make the map more uniform across the field of view. This map is like a correlation map, high in areas in which pixels are correlated to each other, and is saved in the detection output as ‘Vcorr’. To estimate a threshold on the map values for ROI seeds, we find the local peaks in this map by maximum filtering the map with a 3×3 filter size, and finding all pixels with original values greater than their maximum filtered values. The threshold for the ROI seeds is set as `threshold_scaling` times the median of the peak values.

We next iterate over the maximum local values of the map in groups of 200 ROIs, before recomputing the map. We take the largest value in the map as a seed for the ROI and define its initial PC values (**v**) as *U*_smooth_ at the seed. The pixel weights ‘lam’ (λ) are defined as the dot product between *U*_cell_ and **v** for each pixel. We iteratively extend the ROI by one pixel on each edge; pixels are removed if they are less than one-fifth of the maximum. The pixel weights are normalized to be unit norm. After each extension, in this first phase, we recompute **v** as *U*_cell_λ. This extension procedure is repeated until no more pixels are added to the ROI. We subtract the predicted ROI values for each pixel λ**v**^⊤^ from *U*_cell_, and we set the ROI pixel values (with one surrounding pixel added) in the map to zero. We perform this procedure up to 200 times, finding the next largest value in the map, as long as this value is greater than the threshold.

After each group of ROI detections, we recompute the neuropil PC values *N* and ROI PC values *V* . We solve *U* = Λ*V* + *SN* via linear regression with an L2 regularizer of size 1e-3. We then update *U*_cell_:

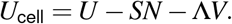

With this updated *U*_cell_ and *V*, for each ROI we refine its pixel weights and extend the pixels with the same procedure as above.

ROI detection continues in groups of 200 until the maximum number of iterations `max_iterations` are reached, or there are no more ROIs found. Two more iterations of optimization are added at the end, where the temporal components of the cells are kept fixed, while the spatial components are updated using the un-smoothed PCs of the movie (if parameter `smooth_masks` is False). This step ensures that the ROI masks do not get over-smoothed. Before these two iterations, the ROIs are reduced to their connected regions if the parameter `connected` is True. The neuropil timecourses are still updated during this step, since the neuropil masks are kept fixed.

#### Cellpose algorithm

The user can alternatively run an anatomical algorithm to find ROIs. For this we provide Cellpose, a generalist cellular segmentation algorithm [39]. The image used for segmentation is specified by setting `img`, and can be the ratio of the maximum projection image and the mean image (‘max_proj / meanImg’), the mean image (‘meanImg’), or the maximum projection image (‘max_proj’). This image can be high-pass filtered before running Cellpose, with setting `highpass_spatial`. The user can specify the Cellpose model with setting `cellpose_model`.

The pixel weights in each ROI ‘lam’ are the pixel values from the maximum projection image if ‘max_proj / meanImg’ or ‘max_proj’ are used as the segmentation. If the mean image is used, then the weights are the mean image clipped at the first and 99th percentile and scaled between 0.1 and 1.1. The ‘med’ center of the ROI is the median of the pixels in the ROI, or the pixel within the ROI closest to the median if the median is outside the ROI.

### Signal extraction

Next we extract the time course for each ROI and the corresponding neuropil signal for each ROI. The settings for signal extraction are contained in the dictionary ‘extraction’. ROIs can be overlapping (see example in Figure 4f). By default we do not use pixels that are in multiple ROIs for signal extraction,but this can be turned off by setting `allow_overlap` to False. The ROI mask values are defined using the ‘lam’ weights for each pixel, normalized to sum to 1 for each ROI.

The neuropil is the signal from the dendrites and axons surrounding the ROI that are captured within its point spread function, and it can be estimated by using the activity surrounding the ROI [71]. We define the neuropil masks as squares around each ROI, with a minimum size `min_neuropil_pixels` and with a padding around the ROI of `inner_neuropil_radius` in pixels. Other ROI pixels are excluded from the neuropil masks. In fields of view with many ROIs, this may exclude many pixels and reduce the accuracy of the neuropil estimate. Thus we do not exclude all ROI pixels and only exclude pixels with sufficiently high ‘lam’ weights, determined by an adaptive percentile threshold. The percentile filter is applied in a window of five times the median ROI radius with a percentile set by `lam_percentile`, and the pixels less than the filter output are not excluded from the neuropil estimation. The neuropil mask weights are defined to be equal in size, as one over the number of pixels in the mask.

To extract the ROI and neuropil traces, a matrix multiplication is performed between the ROI and neuropil masks and the registered frames, in batches defined by the setting `batch_size`. This matrix multiplication is a sparse matrix multiplication that is accelerated on the GPU. The outputs are defined as ‘F’ and ‘Fneu’ for the ROI and neuropil traces respectively. The default neuropil coefficient is 0.7, and is specified by the setting `neuropil_coefficient`. Before spike deconvolution, the neuropil trace is subtracted from the ROI trace with the neuropil coefficient. Then an additive baseline is estimated for each trace. The settings for the baselining are in the dictionary ‘dcnv_preprocess’. First the trace is smoothed by a Gaussian filter of standard deviation `sig_baseline` frames. In the default baseline computation, ‘maximin’, the rolling maximum of the rolling minimum is computed in a window of `win_baseline` seconds. This baseline is then subtracted from the trace, and then the trace is deconvolved using OASIS with no sparsity constraints [12]. OASIS requires the input of the timescale of the indicator decay, which is defined in the pipeline as `tau`* `fs`. We accelerated the OASIS deconvolution using numba, in batches of frames of size `batch_size` [55].

### ROI statistics and classification

We compute several statistics of the ROI shape and the neuropil-corrected fluorescence trace, which are saved in the dictionary for each ROI, which contains ‘ypix’, ‘xpix’, ‘lam’ and ‘npix’ as defined during ROI extraction:

- ‘soma_crop’: We estimate the pixels in the somatic compartment of the ROI by computing the sum of pixel weights (‘lam’) within a radius of the center ‘med’ of the ROI. We compute the difference of this sum as a function of the radius. The radius at which this difference is less than one-third of its maximum value is defined as the radius of the somatic compartment of the ROI, if this radius exists - otherwise all pixels are considered to be somatic. The ‘soma_crop’ key is a boolean list with True values for the pixels in the somatic compartment. The total number of soma pixels is saved in ‘npix_soma’. Note: for all the following shape estimates, only the somatic pixels are used - the user can turn off the somatic cropping by setting `soma_crop` to False in the ‘detection’ settings.
- ‘compact’: We compute how compact an ROI is, where a compactness of 1.0 is equivalent to a disk, and larger values mean the ROI is less compact than a disk. We compute the median pixel of ‘ypix’ and ‘xpix’, and compute the mean Euclidean distance of each pixel from this median (saved as ‘mrs’). This disk normalized by the mean Euclidean distance for the pixels in a disk ROI with the same number of pixels (saved as ‘mrs0’).
- ‘radius’: The radius of the ROI, estimated as two times the major axis radius of a multivariate Gaussian fit to the pixel weights.
- ‘aspect_ratio’: The ratio between two times the major axis radius and the sum of the major and minor axis radii from the multivariate Gaussian fit. This produces an aspect ratio between 1.0 and 2.0 where larger values indicate more elongated ROIs.
- ‘npix_norm’: The number of pixels ‘npix_soma’ are normalized by the the median size of the first 100 ROIs extracted (largest variance explained), or by the median size of all ROIs if Cellpose is used for extraction. ‘npix_norm_no_crop’ uses ‘npix’ for this computation instead.
- ‘overlap’: boolean array specifying which pixels in the ROI are overlapping with other ROIs.
- ‘snr’: An estimate of the SNR of the activity trace *x*, defined as

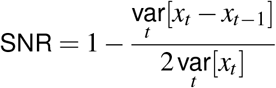

- (similar to [72]).
- ‘skew’: The skewness of the activity trace.
- ‘std’: The standard deviation of the activity trace.

The ROI classifier in Suite2p by default uses the ‘skew’, ‘npix_norm’, and ‘compact’ features. If applied before signal extraction (with `preclassify`), only ‘npix_norm’ and ‘compact’ feature are used. The classifier is trained on ground-truth labels of what is a cell and not a cell (saved in ‘iscell’), which the user can label in the GUI. The classifier is a type of weighted, non-parametric, naive Bayes classifier [73]. Each feature is non-parametrically binned into 100 bins separately for the cell and not-cell classes. The probabilities in each bin are smoothed across bins by a Gaussian with standard deviation of 2 bins. The log posterior likelihood of each feature is then used as input to a logistic regression classifier that is optimized with the ‘liblinear’ solver, using an L2 regularization of size 100 [56]. Effectively, this secondary classification step weighs the importance of each feature relative to the others. A simple naive Bayes classifier would simply use weights of 1 for each feature axis.

### Parameter guidance for different recording types

We always recommend trying the default Sparsery algorithm first, as we have found it can work for many data types and has the flexibility to detect ROIs with various sizes and shapes. For Sparsery detection it is important to ensure that the `highpass_neuropil` scales with the size of the ROI diameter in pixels, with a scaling of approximately 3x the ROI diameter. For recordings with long dendrites, it may be necessary to run the algorithm with a few different parameter values, starting with 3x the dendritic with and increasing. Sourcery and Cellpose detection will only work for round ROIs (somatic recordings), and the `diameter` needs to be set accurately to the diameter of the ROIs in pixels for optimal detection.

For one-photon recordings, motion correction usually requires parameters different from the default. We usually recommend increasing the `spatial_taper` to 100 to reduce the impact of the vignetting of the field of view on the registration. We also recommend increasing the `smooth_sigma` to 3, depending on the zoom this may not need to be as large. For ROI detection, we recommend Sparsery. Because the signals are often slower, we recommend reducing the `highpass_time` parameter to 15 frames or lower. The ROIs are often heavily overlapping so it may be necessary to increase `max_overlap` to 0.85 or higher to reduce the number of ROIs removed due to overlap.

For voltage imaging recordings, we recommend using Cellpose, or fine-tuning a Cellpose model specifically for your data (e.g. in [74]).

For low-SNR or short recordings with somatic ROIs, we recommend trying the Sourcery detection algorithm, with the potential of using the `smooth_masks` parameter to increase ROI pixels for extraction. We also recommend the `denoise` option, which applies PCA denoising to the binned movie in blocks across pixels. If these options do not work, then we recommend using the Cellpose detection algorithm, which can be applied to the mean image, the maximum projection image, or a ratio of these. It may be necessary to fine-tune a Cellpose model for your specific dataset type [75].

### 100,000 neuron recording analysis

For the 100,000 neuron recording, we ran Suite2p with default settings, with `tau` set to 0.25 for jGCaMP8s and the detection threshold set to 0.8 and max ROIs set to 200000 for Sparsery. We kept all ROIs with a cell probability above 0.45.

During the recording, the mouse ran in a closed-loop virtual reality with six corridors (Figure 1d). We computed the tuning curve of each neuron to each corridor by interpolating to 22 equally spaced positions within each corridor, and averaging the responses across odd (training) trials and even (test) trials. This produced two average tuning curves to each corridor for each neuron - a training and test tuning curve. To compute the reliability of the responses of the neurons, we correlated the training and test tuning curves across all corridors and positions for each neuron (length 132 in total) (Figure 1e). We computed the maximum response of the neuron across the training tuning curves to obtain a preferred corridor and preferred position for each neuron. Then we plotted the test tuning curve for each neuron in its preferred corridor sorted by preferred position (Figure 1f). To quantify correlations across planes, we correlated the training tuning curve for each neuron in a plane to the test tuning curve of its closest neighbor in the following plane (Figure S2)d.

### Benchmarking motion correction

To benchmark motion correction, we used a recording in which neurons expressed both a functional and an anatomical marker (GCaMP6s and td-Tomato), recorded in separate imaging channels (Figure 3a). We applied the motion correction algorithms to the recording, using the functional channel to compute the motion shifts and applying these shifts to the anatomical channel. The Suite2p motion correction pipeline was run with the defaults. The Caiman v1.13.0 motion correction was run with the defaults, other than the maximum rigid shifts, `max_shifts`, and the maximum non-rigid shifts,`max_deviation_rigid`, which were set to (65,65) and 10 respectively, which were an upper bound on the deviations in this dataset, and which matched the defaults in Suite2p. We used FIOLA git commit ec7cb37c85831d14fb6d2402ca1aa5316e8ec886. We also updated the Fiola parameter for maximum shifts `ms` to [65, 65], but otherwise kept the parameters as the defaults. The default for initial frames was one tiff, which consisted of 500 frames in this case. We computed the registration metrics for the functional and anatomical channels after registration with each algorithm (Figure 3b-f).

For quantifying timing performance, we ran each algorithm on a subset of the tiffs, from 500 to 32,000 frames (Figure 3g). The Caiman algorithm was run on 16 cores on an Emerald Rapids compute node. The Suite2p and Fiola algorithms were run on 12 cores on a Cascade Lake compute node with an A100 GPU. The frame processing rate in Hz was estimated as the slope of the number of frames as a function of the runtime.

### Benchmarking detection

#### Hybrid ground-truth generation

To benchmark detection, we generated hybrid ground-truth recordings (Figure 5a). For this, we recorded three mice expressing RiboL1-jGCaMP8s so that there were no background neuropil signals [37]. We computed the maximum projection image (maximum across frames in time) for each recorded plane, and extracted the “ground-truth” ROIs from this image using the Cellpose SAM model [76]. The Cellpose SAM model was run with settings `diameter`=16, `cellprob_threshold`=-3, and `flow_threshold`=0. From each detected ROI, the fluorescence trace for the ROI and the neuropil signal were extracted using default Suite2p settings. The neuropil-corrected traces were then used as the ground-truth traces (the neuropil correction is performed to remove frame-wide changes in brightness).

For the hybrid ground-truth recording, we downsampled the recording by a factor of two spatially, so that the pixel resolution was 0.5 *µ*m, more similar to many imaging setups. This downsampled recording is the base onto which we added simulated axons/dendrites, simulated neuropil, and shot noise. Accordingly, the ground-truth ROIs were also downsampled spatially by a factor of two for comparison with the detected ROIs.

For the simulated axons/dendrites and simulated neuropil, we removed slow fluctuations in time from the recording. This was performed by estimating a per pixel baseline by maximum filtering of a minimum filter of the recording, with a filter width of 1 second, and then subtracting this baseline from each pixel.

After this filtering, we extracted fluorescence traces for each Cellpose ROI, which were used for the axons/dendrites. For the simulated neuropil, we downsampled the filtered recordings by a factor of two spatially, and also subtracted the per pixel mean from the recording, using the mean pixel intensities from the first 2,000 frames.

The simulated axon/dendrite ROIs were ellipses with centers drawn from a uniform random distribution across the field of view (*x*_0_, *y*_0_) - the default number of ellipses was set to 2,000. The ellipse radii, for one axis of the ellipse, were drawn from a uniform distribution between 0.75 and 2. For the radii of the other axis, two-thirds of the ellipses had radii drawn from a uniform distribution between 0.75 and 2, and one-third had radii drawn from a uniform distribution between 1 and 6. For the pixel weights ‘lam’ of each ellipse ROI, we added Gaussian random noise to the output of the ellipse equation for each pixel, 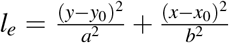,and rectified the values (set the minimum to 0). The ‘lam’ pixel weights were then set to 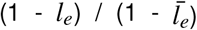, so that the largest weights were at the center of the ROI, and that the weights summed to the number of pixels in the ROI. We varied the number of such ROIs (Figure 5e-g left), keeping the other parameters at their default values. For the fluorescence traces, we sampled activity traces with an SNR *>* 0.3 which were nearby spatially on the other three planes: we computed the Euclidean distance between the ellipse ROI center and the ROIs on the other planes, and used the activity trace from the *n*^th^ closest ROI, where *n* was drawn from a random exponential of mean five and rounded to an integer. These activity traces were added to the recording, weighted per pixel by the ‘lam’ weights of the ROI.

The simulated neuropil consisted of 7×7 basis functions in X and Y, defined as the outer product of sines and cosines, with the frequencies from one to three cycles across the field of view. Each basis function was normalized to be unit norm. Then the activity of each basis function was simulated by projecting the basis function onto the filtered and downsampled recordings from the other three planes, summed across the planes. Then this activity was projected onto the basis functions to create the simulated neuropil. After projection, the per pixel mean from the downsampled recording without filtering was added, and then the neuropil activity was rectified (minimum set to 0). Then the neuropil activity was scaled by the neuropil fraction (default 0.4) and added to the recording, which was scaled by one minus the neuropil fraction. We varied the neuropil fraction (Figure 5e-g middle), while keeping the other parameters at their default values.

Next we simulated shot noise using a Poisson distribution, with a scaling term, the Poisson noise scale, which was set to 20 by default. The recording was divided by the Poisson noise scale then used as the mean of a Poisson distribution, which we sampled from to generate the intensity of each pixel across all timepoints in the recording. The output was then multiplied by the Poisson noise scale to ensure the mean pixel intensity was similar across scales. We varied the Poisson noise scale (Figure 5e-g right), while keeping the other parameters at their default values.

#### Performance metrics

We compared the ground-truth ROIs and the detected ROIs, using their pixel distributions and their activity traces. We excluded ROIs with an SNR less than 0.25, assuming these may be non-active or noisy ROIs detected by Cellpose or the other algorithms (Figure 5b). We also applied a size filter on the ground-truth and detected ROIs, excluding ROIs smaller than the fifth percentile or larger than the 95th percentile of ground-truth ROI sizes.

We computed the intersection-over-union (IoU) of the detected ROIs and the ground-truth ROIs, which is the number of overlapping pixels divided by the number of unique pixels across both ROIs. For Caiman, there were several pixels with small weights, and thus we thresholded the detected ROI pixels such that only pixels above 0.1 times the maximum weight of the ROI were kept. If the IoU value was larger than 0.5, then the ROIs were considered to be matched. We next filtered the matched ROIs so that each ground-truth ROI could only be matched to one detected ROI. The remaining matched ROIs were defined as the “true positives” (TP). The “false positives” (FP) were the detected ROIs without matched ground-truth ROIs, and the “false negatives” (FN) were the ground-truth ROIs without matched detected ROIs. The numbers of TP, FP, and FN ROIs were computed for each plane in each recording, and summarized by the F1 score: 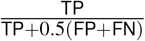

#### Detection algorithms

Suite2p Sparsery and Sourcery were run with all default settings, with `tau` set to 0.25 because the indicator was jGCaMP8s. We performed a sweep across the `threshold_scaling` parameter, from 0.6 to 1.4, to determine the optimal threshold for ROI detection, defined by F1 performance averaged across all recordings (Figure S6ab). The optimal threshold was determined to be 0.7 for Sparsery and 1.0 for Sourcery, and this was used for all benchmarking in Figure 5.

Caiman v1.13.0 was run with the settings used for a V1 recording with a 2 *µ*m/pixel spacing from their paper (called ‘yuste_params’). We performed a sweep across the `K` parameter, which specifies the number of components per patch, from 7 to 13 (Figure S6b). The optimal `K` was determined to be 9, and this was used for all benchmarking in Figure 5.

We also performed a grid search over several of the parameters in Caiman, centered at the parameter values used in Figure 5: autoregressive order (p), number of background components (gnb), number of components (K), Gaussian smoothing (gSig), and patch half-size (rf). We did not observe substantial improvement in the performance of Caiman when varying these parameters (average F1 score of 0.44 versus 0.49 for best parameter combination). In Figure S7, we plotted the parameter combination that produced the highest F1 score for each data point.

The Caiman algorithm was run on 16 cores on a compute node, and the Suite2p algorithm was run on 12 cores on a compute node with an A100 GPU.

#### Caiman ground-truth datasets

We compared Suite2p performance to Caiman on datasets which they curated (Figure S8) [31, 50]. Giovannucci et al used subsets of the ground-truth data (in windows of the full images) as part of the training set for their classifier, without holding out any of the datasets, and thus these datasets constituted the training data for their classifier. We did not re-run Caiman, using the saved outputs from the zenodo page.

We ran Suite2p Sparsery and Sourcery with `tau` set to 0.4. The `highpass_neuropil` settings were 40 for J115 and K53, and 25 for YST, because the recordings were at different zoom levels. The diameter for Sourcery was set to 16 for J115 and K53, and 8 for YST given the different zoom levels. The threshold for Sourcery was set to 0.3, and the `max_overlap` set to 0.85. The cell probability thresholds were set to 0.35 for Sparsery and 0.25 for Sourcery. We additionally filtered by size, removing all ROIs with an `npix_norm` less than 0.4.

We evaluated the performance of the algorithms the same way as the hybrid ground-truth datasets, matching the ground-truth and detected ROIs spatially. We used IoU thresholds of 0.2 and 0.5, showing 0.2 to replicate the threshold chosen in the Caiman paper - although this is a very small threshold for matching.

### Cell tracking analysis

For the cell tracking imaging sessions, we ran Suite2p with default settings, with `tau` set to 0.25 and the detection threshold set to 1.0 and max ROIs set to 20000 for Sparsery. We kept all ROIs with a cell probability above 0.5. We ran Suite2p with these settings per session, and also with all sessions together (choosing all the session folders for processing).

We ran Track2p (git commit 9c1fc2aa3acdcf562e3af0b06760f115145685e3) using the single session folders with default settings [45]. The Track2p paper used the correlation of pairwise-correlations between neurons across sessions as a measure of success of tracking. We performed the same analysis, correlating the pairwise-correlation matrices to the first session for each session (Figure 6de).

The mouse ran through two corridors in closed-loop virtual reality, with one corridor delivering rewards. We interpolated the responses of each neuron to 30 positions within each corridor to produce single-neuron tuning curves to each corridor, producing a tuning curve of length 60. We correlated each neuron’s tuning curve in each session to its first session tuning curve, to estimate the consistency of each neuron’s response properties across days (Figure 6fg).

### Neuropil analysis

Using the example recording for motion correction, we estimated neuropil correction coefficients empirically from the ROI and neuropil traces (‘F’ and ‘Fneu’). For this, we applied FastICA from scikit-learn with two components to the ROI and neuropil trace for each ROI [56]. The component with the highest absolute correlation to the neuropil trace was considered to be the neuropil component, and the other component then was defined as the ROI component, multiplied by the sign of the neuropil correlation. The neuropil coefficient was defined as the ratio of the ROI component weights of the neuropil and ROI trace times -1.

### Other imaging-based recordings

We ran Suite2p on a recording with interneurons, a one-photon miniscope recording, and a voltage imaging recording Figure S9).

For the interneuron recording, we ran Suite2p on the functional recording with default settings, with `tau` set to 0.25 for jGCaMP8s and the detection threshold set to 1.0 and max ROIs set to 20000 for Sparsery. We kept all ROIs with a cell probability threshold above 0.05. We used the mean image from the functional recording as the reference image to register the recording of both the green and red channels. The threshold on red cell probability was set to 0.6.

The one-photon miniscope recording was performed at a frame rate of 30Hz in freely-moving rats in the ventral tegmental area (VTA), injected with GCaMP6s [40, 48, 71]. We ran Suite2p with Sparsery with the detection threshold set to 0.75, the `highpass_neuropil` set to 60, and the `max_overlap` set to 0.85. Because it was one-photon imaging, we set the `highpass_time` for detection lower to 15 frames, and the neuropil coefficient to 1.0. For registration, due to the vignetting of the field of view, we increased the `spatial_taper` to 100 and the `smooth_sigma` to 3. With the increased pixel size and large movements, we increased the `block_size` to 128 and the `maxregshiftNR` to 10. For the Caiman analysis of the recording, we ran the ‘demo_pipeline_cnmfE.ipynb’ notebook.

The voltage imaging was performed at a frame rate of 1002Hz in transgenic larval zebrafish expressing Ace-mNeon2-Kv2.1 using a custom oblique plane microscope [42, 49, 77]. We ran Cellpose inside Suite2p on the mean image, using the latest default model (‘cpsam_v2’), `diameter` set to 8, `cellprob_threshold` set to -6, `flow_threshold` set to 0, and `sharpen_radius` set to 5. For the Volpy analysis of the dataset, we ran the ‘demo_pipeline_voltage_imaging.ipynb’ from Caiman [41].

---

**S1:**
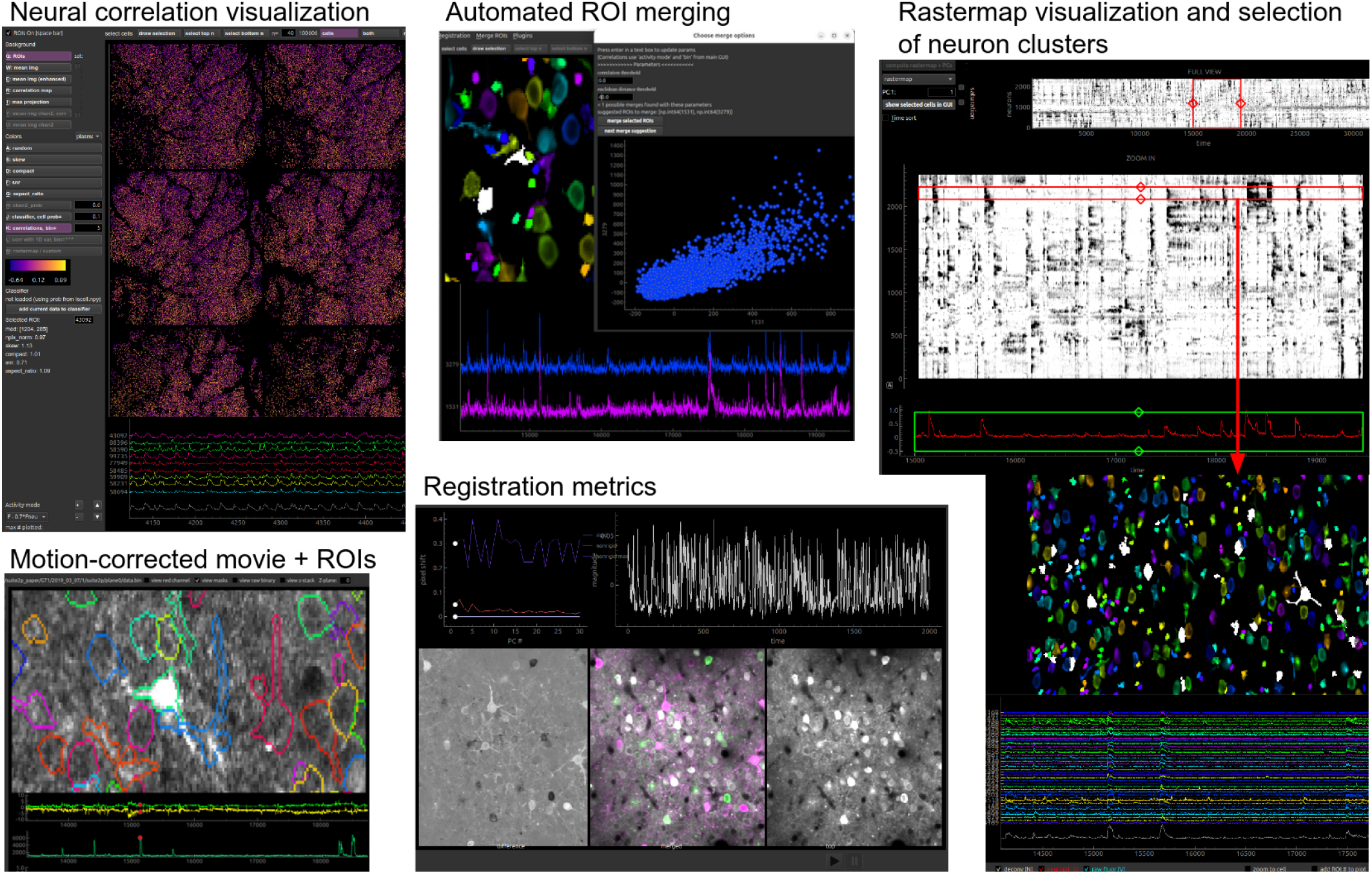
Example functionality of the Suite2p graphical user interface.

**S2:**
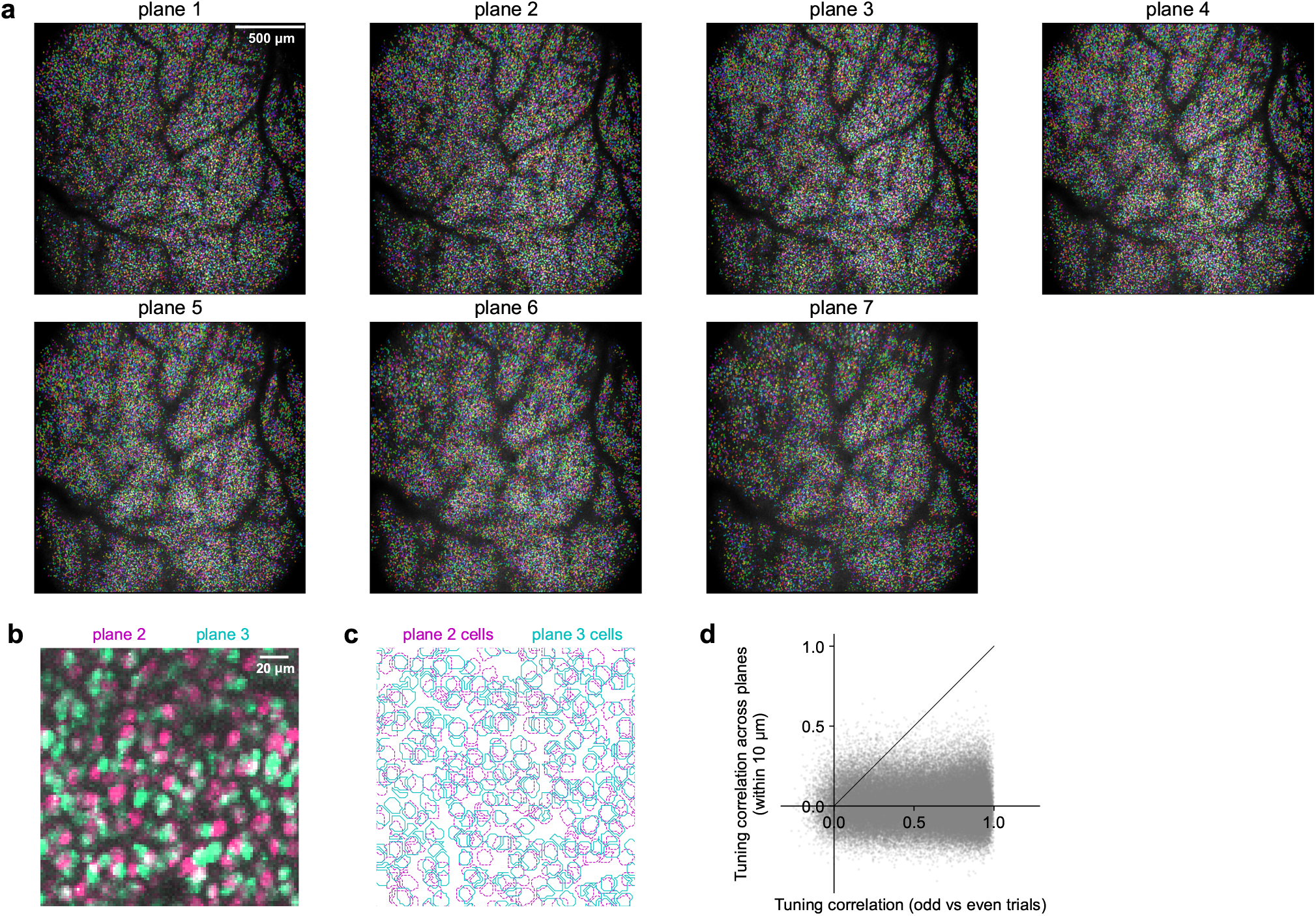
Visualizing ROIs detected in 100,000+ neuron recording. **a**, Each panel shows the maximum projection image (maximum over time) with the detected somatic ROIs overlaid in random colors. Recording was made with a Thorlabs Bergamo2, Toptica Femtofiber Ultra 920 @ 30Mhz and Nikon 10x, 0.5, Plan Apo, Glyc with water immersion in a riboL1-gcamp8s Tigre x Slc17a7 mouse. **b**, Zoom-in of overlay of maximum projection image of plane 2 and plane 3. **c**, ROIs detected in plane 2 and plane 3. **d**, Scatter plot between tuning correlation (like in Figure 1i) and tuning correlation between a cell and the closest cell in the next plane.

**S3:**
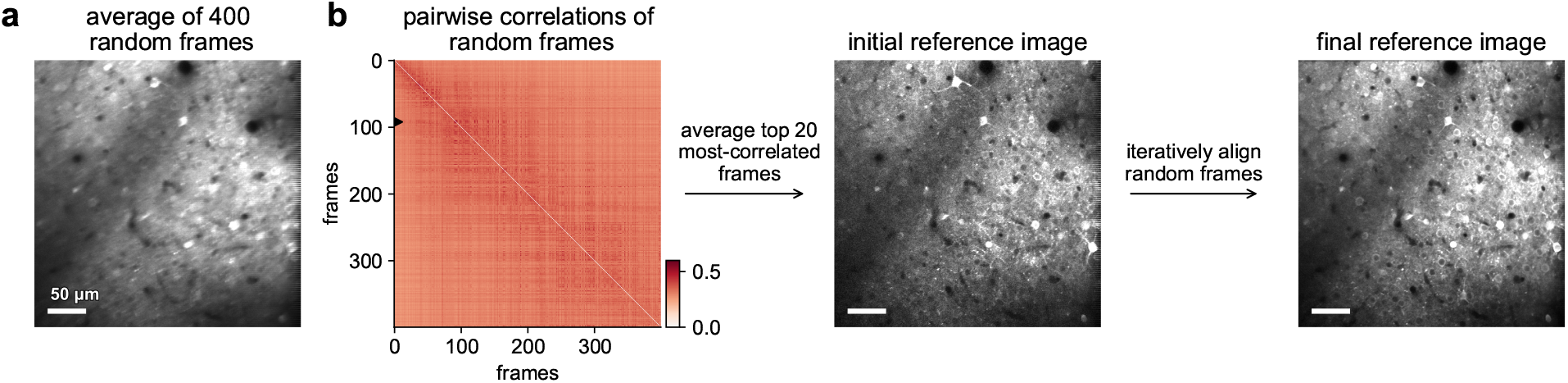
Reference image computation. **a**, Average of 400 un-registered frames. **b**, Procedure to obtain the reference image by finding correlated frames and iteratively aligning them to an increasingly sharper reference image. Left: pairwise correlations of the random frames, with the frame row with the top 20 most-correlated pairs denoted by the black triangle. Middle: Average of 20 most-correlated frames. Right: Average of random frames after iterative alignment.

**S4:**
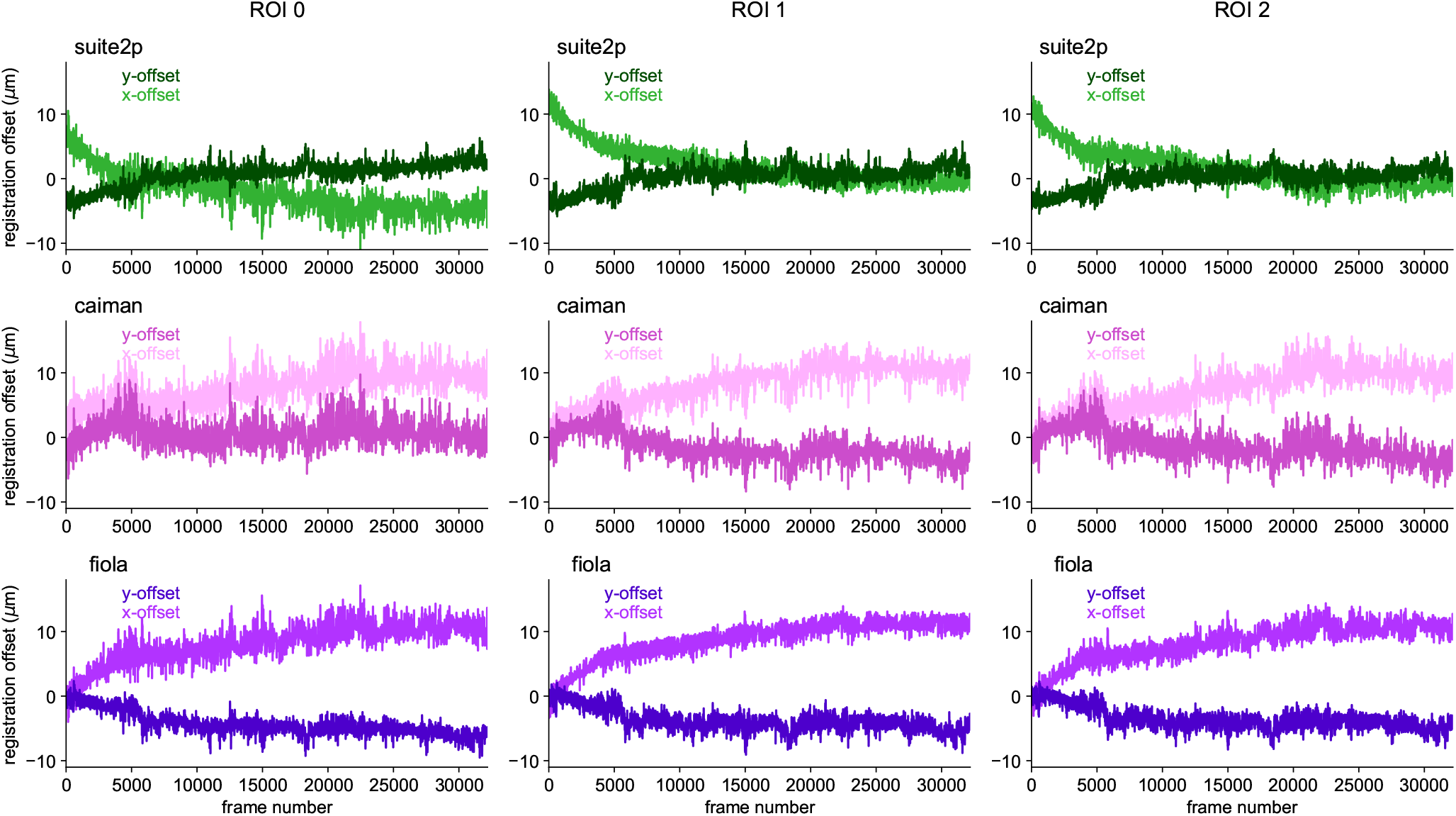
Registration offsets from the experiment in Figure 3. Compared to Suite2p and Fiola, the registration offsets in Caiman span a shorter range, likely due to the continuously changing reference frame.

**S5:**
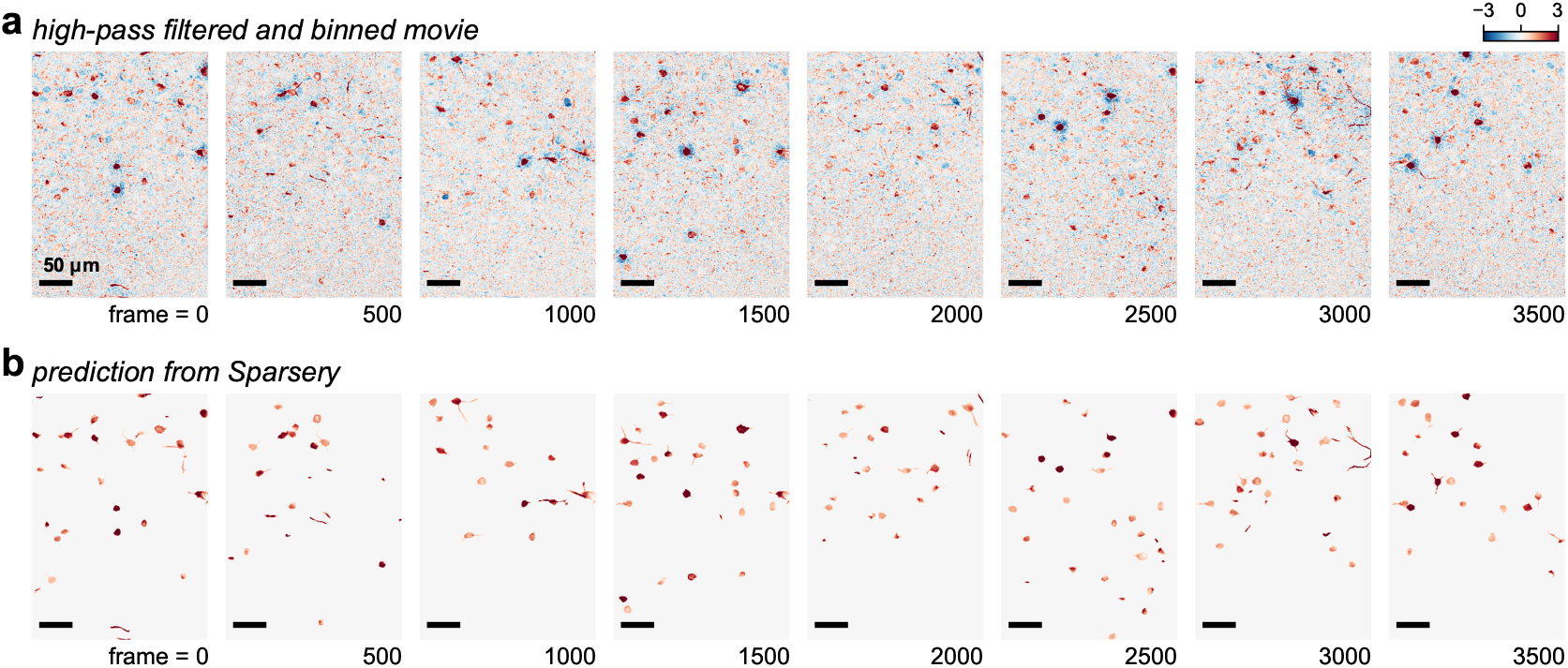
Reconstruction of the movie with Sparsery. **a**, Example frames from the high-pass filtered and binned movie. **b**, Predicted activity from the detected ROIs from Sparsery. Note only positive deviations are reconstructed.

**S6:**
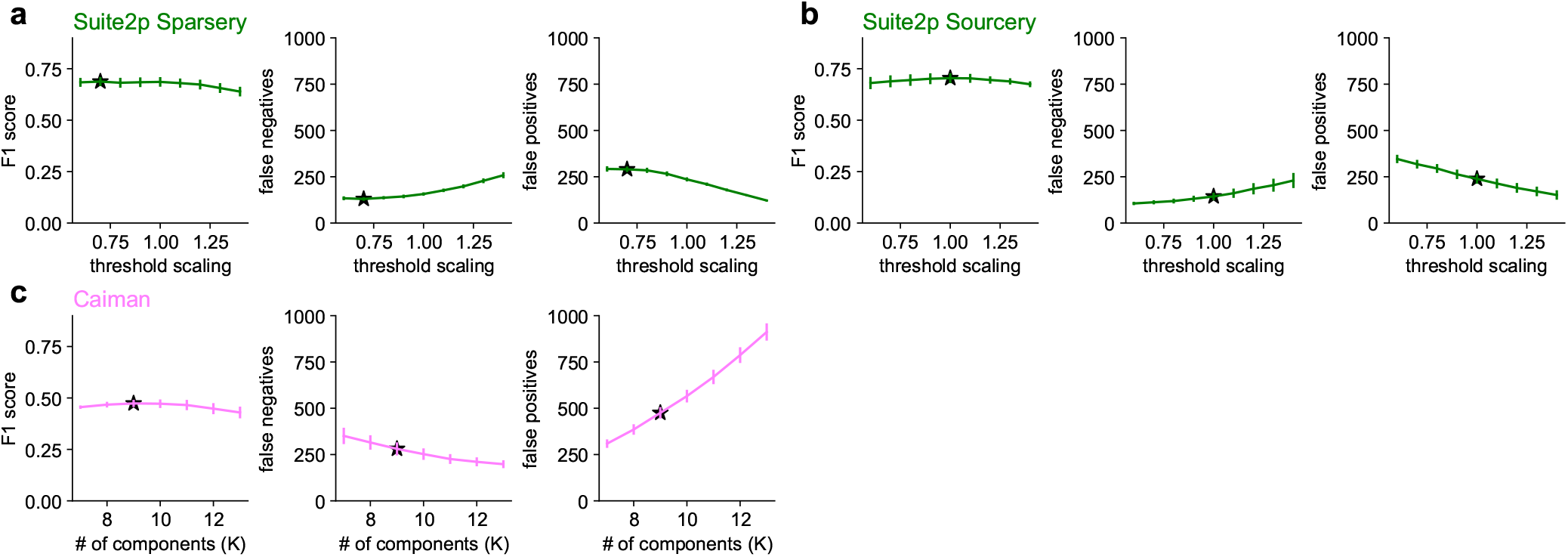
Optimal cell detection thresholds. a, Suite2p Sparsery, **b**, Suite2p Sourcery, **c**, Caiman.

**S7:**
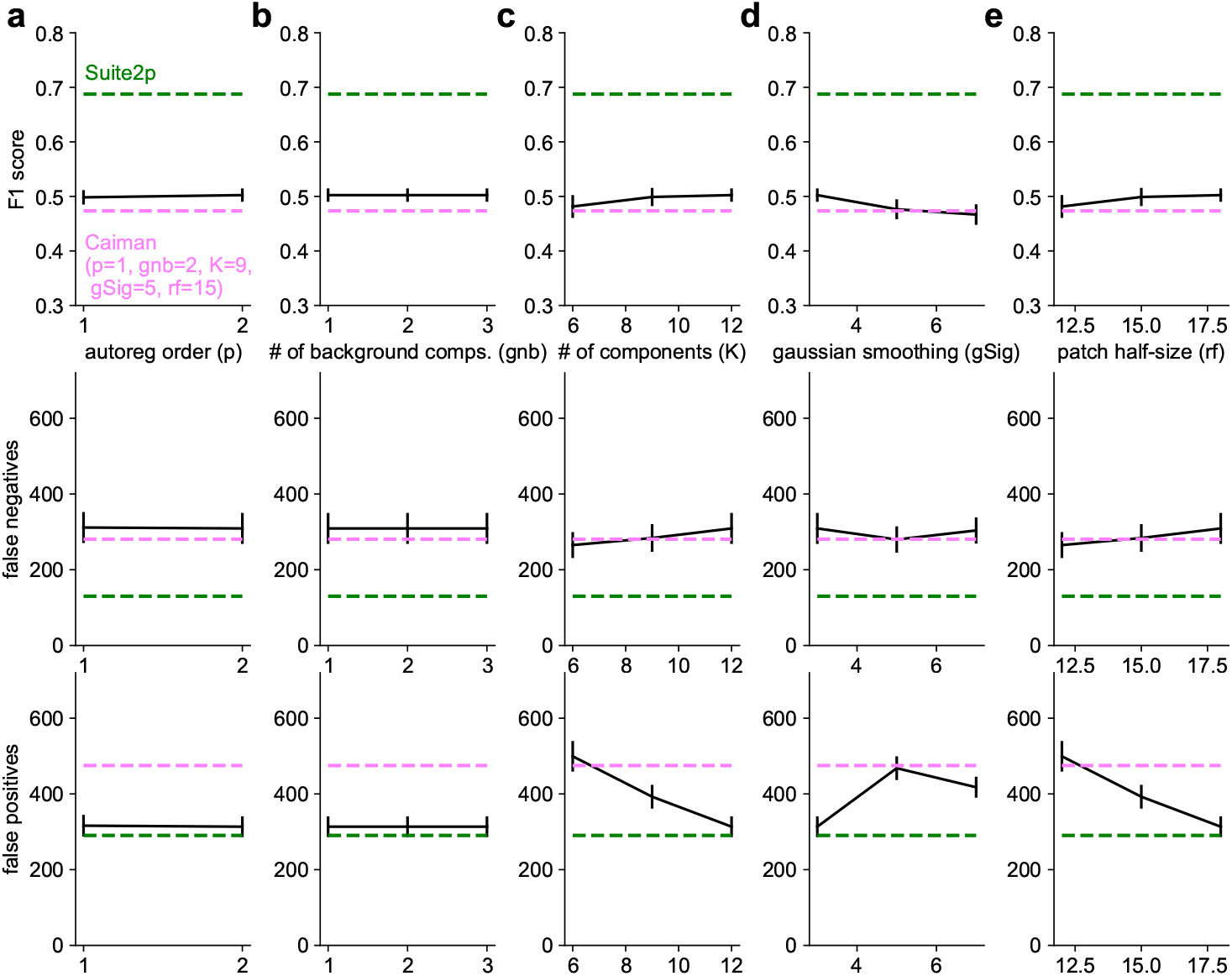
Parameter sweep for Caiman. Each panel shows the detection performance metric (F1 score) as a function of a parameter, after taking the maximum across all other parameters. While this procedure in general overfits, the improvements are minimal regardless. The parameters sweeped over are: **a**, Autoregressive model order. **b**, Number of components for modelling the background / neuropil per patch. **c**, Number of components per patch. **d**, Gaussian smoothing parameter. **e**, Patch half-size in pixels.

**S8:**
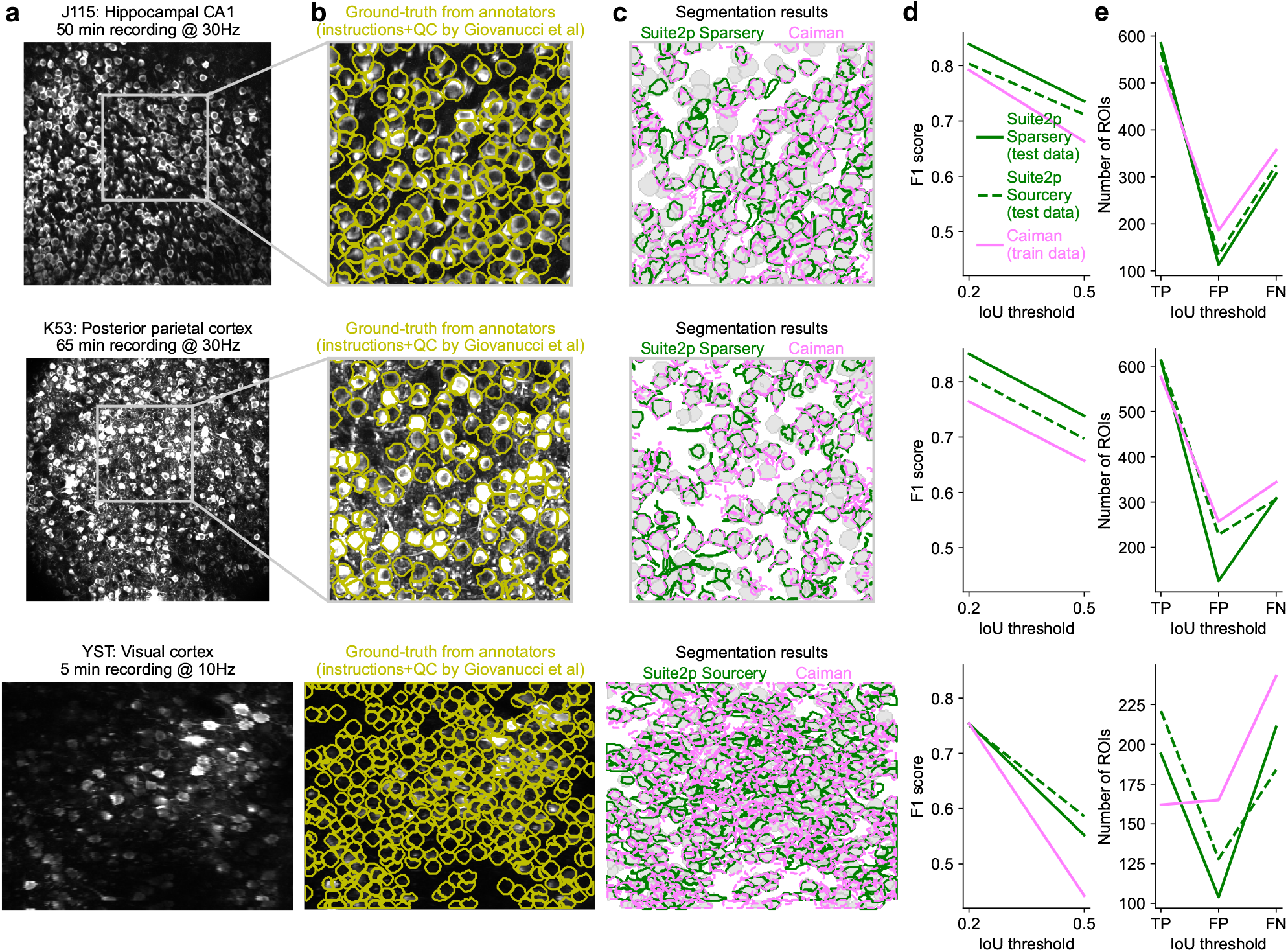
Evaluation on Caiman training datasets. Each row shows a dataset from [31]; there are no scale bars because pixel resolution was not provided. **a** Complete field of view for annotated datasets from [31]. **b**, Ground truth from annotators overlaid on a zoomed in portion of the FOV. **c**, Segmentation results of Suite2p Sparsery and Caiman overlaid on the annotator segmentations. Note that Caiman was trained on the annotator segmentations. **de**, Quantification of match between annotator segmentations and algorithm results, as a function of the IoU threshold used for matching. The very low threshold of 0.2 was kept for comparison with [31], but we note that this can introduce artifacts since very little of the cell masks need to match at this IoU, and unique matches are not guaranteed, unlike the 0.5 IoU threshold. The F1 score is shown in **d**, which is based on the TP, FP and FN numbers in **e**.

**S9:**
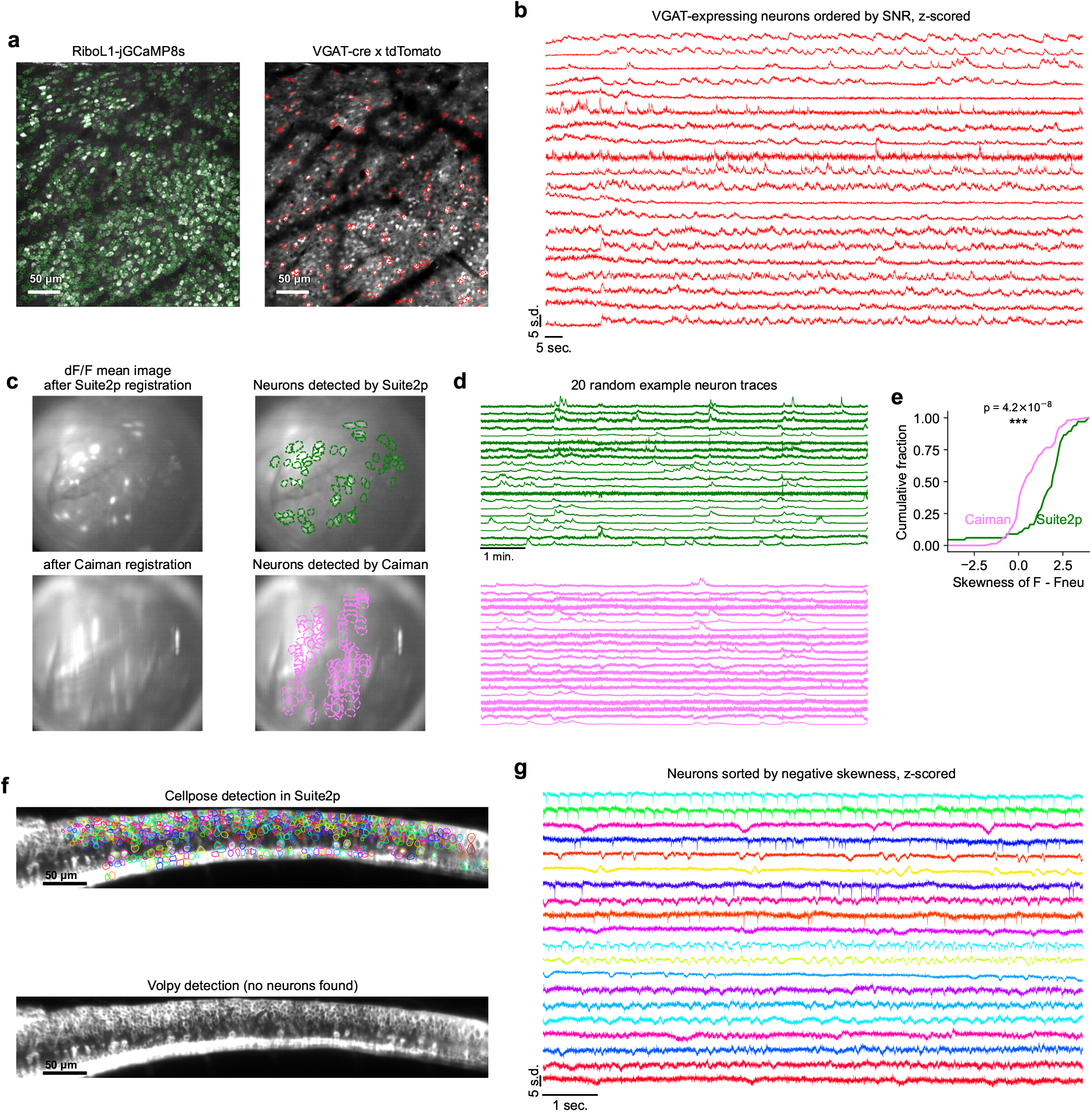
Other imaging-based recordings. **a** (left) Masks detected by Suite2p Sparsery on the green channel of a recording. (right) Subset of the masks on the left, which have a substantially higher intensity of red labeling within vs outside the mask, where the red represents the tdomato channel marking VGAT-positive cells. **b**, Activity of a subset of VGAT-expressing neurons. **c**, Field of view from a miniscope recording using one-photon calcium imaging from [40] (no pixel resolution provided). **d**, Activity of example random neuron traces of masks detected by Suite2p and Caiman. **e**, Cumulative distribution of activity trace skewness of cells detected by Suite2p vs Caiman. **f**, Detection of ROIs in a one-photon voltage imaging application from [42] by Suite2p using the Cellpose algorithm (top) and by Volpy (bottom). Note that Volpy fails here because it expects nuclear labeling [41]. **g**, Example activity traces extracted by Suite2p.

**S10:**
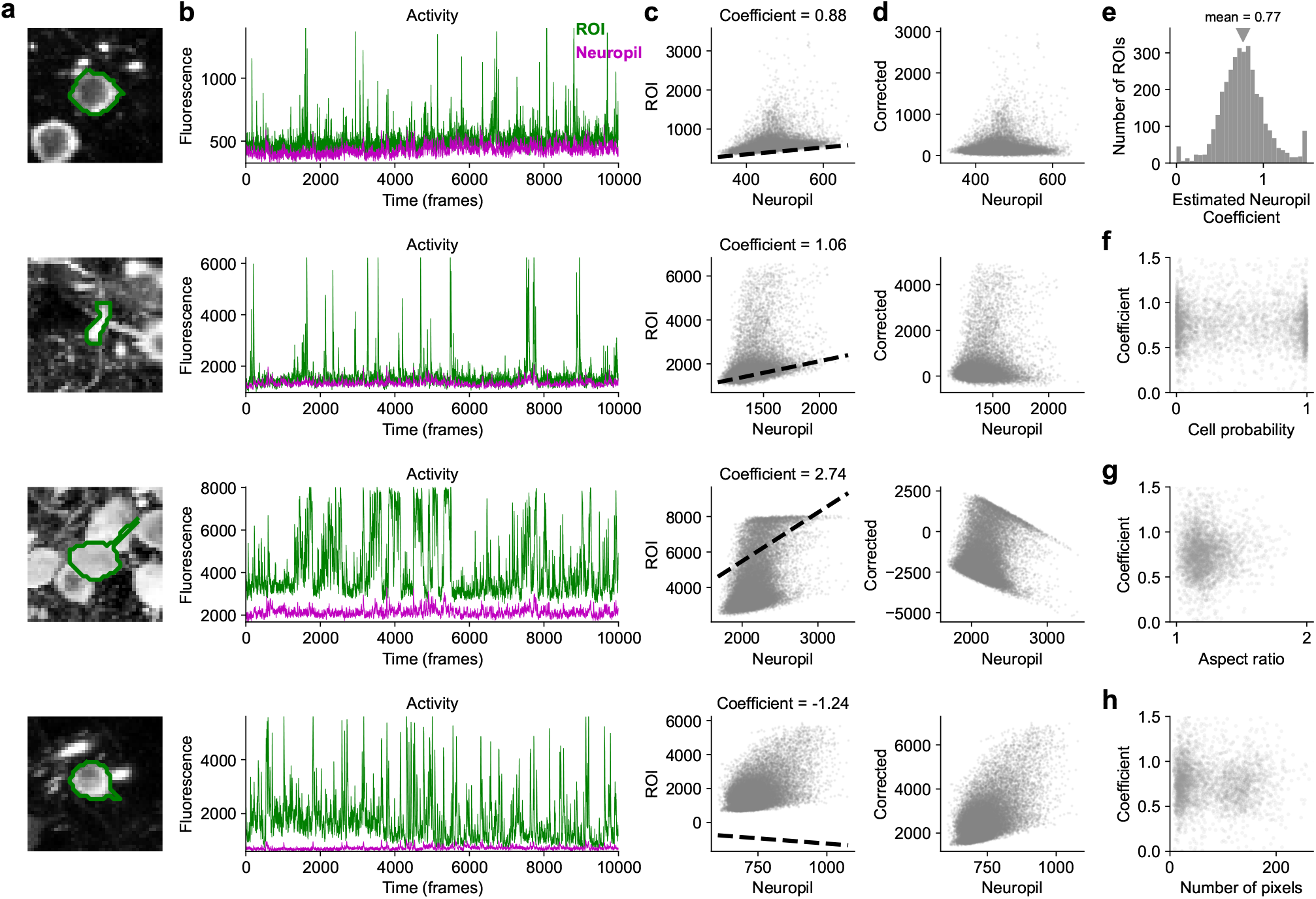
Automatic estimation of neuropil coefficient. Each row is a different ROI. **a** Maximum projection image centered on an ROI detected by Suite2p, area shown is 40 *µ*m x 40 *µ*m. **b**, Activity traces extracted as the average of pixels inside the ROI as well as outside, excluding other cells (neuropil), shown for a subset of the frames. **c**, Scatterplot of activity traces between ROI and neuropil. Fitted dotted line indicates the estimate of the neuropil contamination inside an ROI based on an ICA algorithm. This estimate is consistent for a majority of neurons in this recording, but fails clearly for some ROIs, when the coefficient is either below 0 or above 1, which are not realistic values. **d**, Scatterplot of corrected vs neuropil traces. **e**, Histogram of estimated neuropil coefficients, clipped at zero and 1.5. **f-h**, Scatterplots of estimated neuropil coefficient versus **f** cell probability assigned by the Suite2p soma classifier, **g** aspect ratio, **h** number of pixels in an ROI.

